# Loss of TMEM65 causes mitochondrial disease mediated by mitochondrial calcium

**DOI:** 10.1101/2022.08.02.502535

**Authors:** Yingfan Zhang, Laura Reyes, Junhui Sun, Chengyu Liu, Danielle Springer, Audrey Noguchi, Angel M. Aponte, Jeeva Munasinghe, Raul Covian, Elizabeth Murphy, Brian Glancy

## Abstract

Transmembrane protein 65 (TMEM65) depletion in a patient carrying a homozygous variant in the Tmem65 splice site resulted in severe mitochondrial encephalomyopathy, indicating the clinical importance of TMEM65. However, the function of TMEM65 remains unknown. Here, we generated a TMEM65 reporter mouse as well as whole-body and tissue-specific *Tmem65* knockout (KO) mice to investigate the localization and function of TMEM65. We show that TMEM65 is localized to mitochondria in heart, skeletal muscle, and throughout the brain. Both whole-body and nervous system-specific *Tmem65* KO result in severe growth retardation and sudden death following seizures ~3 weeks after birth, indicating TMEM65 is indispensable for normal brain function. In addition, we find that skeletal muscle-specific *Tmem65* KO leads to progressive, adult-onset myopathy preceded by elevated mitochondrial calcium levels despite unaltered expression of known mitochondrial or cellular calcium handling proteins. Consistently, we demonstrate that ablation of TMEM65 results in a loss of sodium-dependent mitochondrial calcium export. Finally, we show that blocking mitochondrial calcium entry through removal of the mitochondrial calcium uniporter (MCU) rescues the early lethality of whole-body TMEM65 ablation. Our data not only reveal the essential role of TMEM65 in mammalian physiology, but also suggest modulating mitochondrial calcium may offer a potential therapeutical approach to address defects associated with TMEM65 misexpression.

---

TMEM65 was first reported in MitoCarta, an inventory of more than 1000 mammalian mitochondrial proteins^2^, and initially considered to be localized to the inner mitochondrial membrane with three putative transmembrane domains, an N-terminus facing to the mitochondrial matrix, and a C-terminus in the mitochondrial intermembrane space^3^. TMEM65 is encoded by nuclear DNA, and contains a mitochondrial targeting sequence at the N-terminus of the protein so that it can be imported into mitochondria^3^. The expression of TMEM65 is regulated by the long non-coding RNA Steroid Receptor RNA Activator (SRA) ^4^ which forms a ribonucleoprotein complex with leucine-rich pentatricopeptide repeat-containing protein (LRPPRC) and the Stem-Loop-Interacting RNA-binding Protein (SLIRP)^5^. Despite the abundance of information available surrounding the structure and regulation of this protein, the specific function of TMEM65 remains elusive.

Interestingly, TMEM65 has also been suggested to be a plasma membrane protein localized to the intercalated discs in cardiac myocytes where it is proposed to function as a novel regulator that interacts with connexin 43 (CX43) to modulate gap junction communication^6,7^. However, a critical role for TMEM65 in cardiac function was not supported by a recent report of a patient with severe mitochondrial encephalomyopathy caused by a significant loss in TMEM65 expression, as the patient did not show any evidence of cardiomyopathy^1^. Abnormal expression of TMEM65 has also been linked to Barth Syndrome^8^, Covid-19^9^, and several cancers^10-13^ further suggesting a better understanding of the specific function of TMEM65 may be of critical clinical importance.

## TMEM65 KO mice have mitochondrial disease

To investigate the tissue and subcellular expression of TMEM65 protein, we generated a TMEM65-V5 in frame knock-in mouse model wherein a V5 tag was inserted to exon7 of Tmem65 before the stop codon so that endogenous TMEM65-V5 instead of TMEM65 was expressed (*Tmem65^+V5/+V5^*, Fig. 1a). Western blots of TMEM65-V5 mouse tissues indicated TMEM65 expression throughout the body with high expression in brain, heart, and skeletal muscle (Extended Data Fig. 1a). Immunofluorescent analyses of TMEM65-V5 mouse tissue sections confirmed TMEM65 is localized primarily to mitochondria in brain (Fig. 1b and Extended Data Fig. 1b) and heart (Extended Data Fig. 2a-d). In wild type mouse skeletal muscle, TMEM65 was expressed within the grid-like mitochondrial network^14^ surrounding the contractile machinery in both fixed, isolated soleus fibers (Fig. 1c) and in TMEM65-GFP transfected Tibialis anterior (TA) muscle in vivo (Extended Data Fig. 1c). Therefore, TMEM65 is a mitochondrial protein highly expressed in excitable tissues.

**Figure 1.**
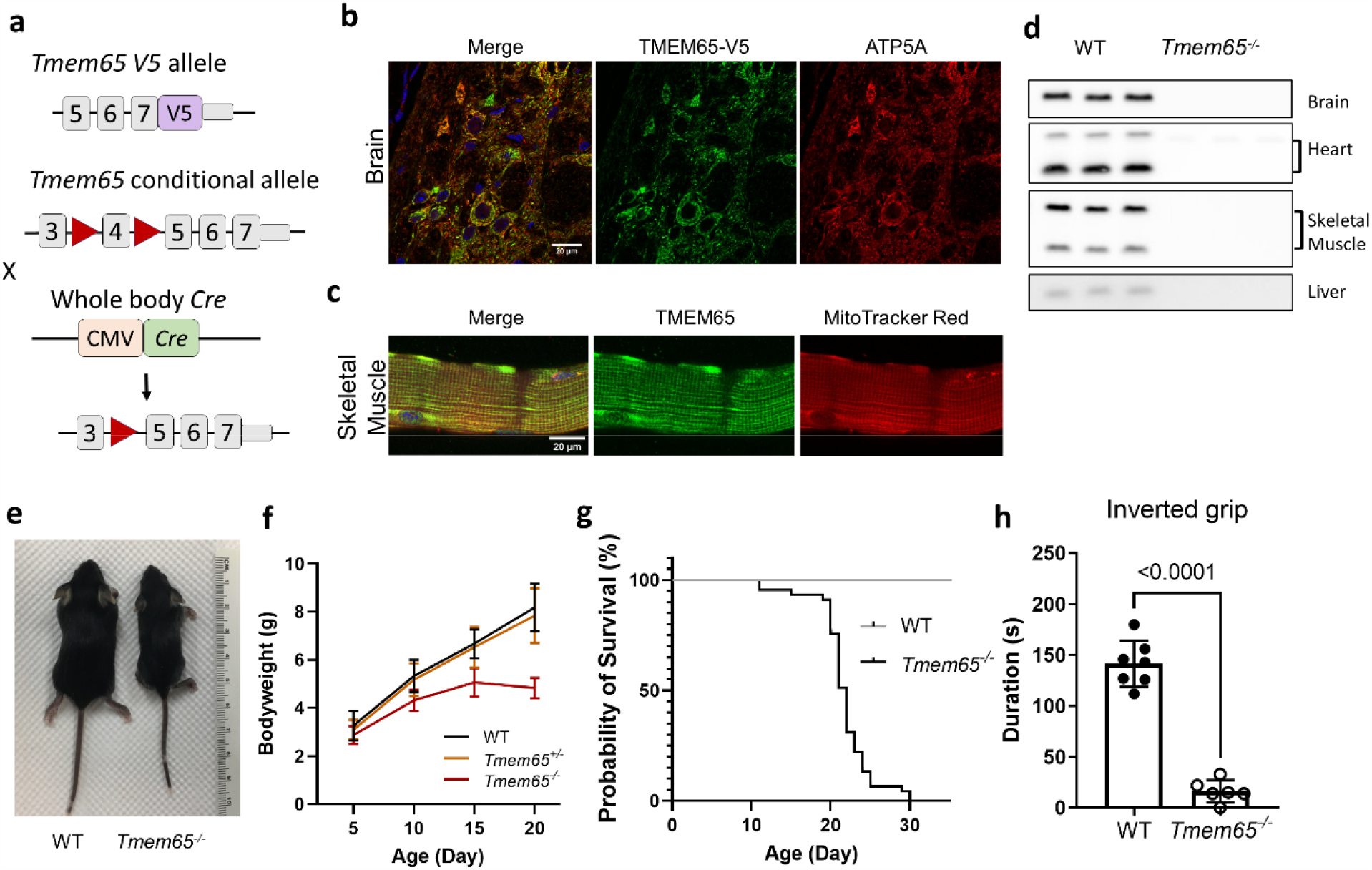
TMEM65 is a mitochondrial protein, and ablation of TMEM65 is lethal in mice. **a**, Schematic of Tmem65-V5 mouse, floxed mouse, and conditional KO mutant mice strategy. In *Tmem65^+V/+V^* mice, V5 tag (purple rectangle) was inserted before the stop codon in exon 7. In *Tmem65^fl/fl^* mice LoxP sites(red triangles) flank exon 4. *Tmem65^fl/fl^* were crossed with whole body Cre (CMV promoter) to generate Tmem65^−/-^. **b**, TMEM65 colocalized with ATP5A, a mitochondrial complex V subunit in mouse brain section near lateral reticular nucleus. **c**, TMEM65 colocalized with MitoTracker Red in isolated mouse soleus skeletal muscle. **d**, Western blotting experiments show loss of TMEM65 in different tissues from P19 Tmem65^-/-^ mice. n = 3 replicates per group. **e**, Representative image of a WT control and Tmem65^-/-^ littermates at P20. **f**, Bodyweights of littermates measured at P5, P10, P15 and P20. WT (Tmem65^+/+^) n = 29; Tmem65^+/-^ n = 48; Tmem65^-/-^ n = 22-28. All data are presented as mean ± SD. **g**, Survival chart of WT controls and Tmem65^-/-^ littermate shows that Tmem65^-/-^ pups (n = 45) died between P11 and P30, with an average life span of 21.7 ± 3.6 days. **h**, Inverted grip test was done on weaning day of P21. WT n = 7; Tmem65^-/-^ n = 6. Individual values as well as mean ± SD are presented.

To investigate the functional impact of TMEM65 in mice, we initially generated global TMEM65 KO mice with sgRNAs targeting exon1 and/or exon4 of Tmem65 using standard CRISPR methodology (Extended Data Figure 3). However, all resultant KO mice died at 2-4 weeks of age precluding breeding. As a result, we generated a conditional loss-of-function mouse model wherein exon 4 of Tmem65 was floxed by LoxP sites (*Tmem65^fl/fl^*, Fig. 1a) and we crossed these floxed mice with the CMV-Cre line^15^ to recreate TMEM65 whole-body KO mice (Fig. 1d). TMEM65 KO mice (*Tmem65^-/-^*) were born at expected Mendelian ratios but were growth retarded and showed growth regression after postnatal day (P) 15 (Fig. 1e,f). All *Tmem65^-/-^* mice died between P11 and P30 with average life span of 21.7 ± 3.6 days (Fig. 1g) recapitulating the initial global TMEM65 KO phenotype. Epilepsy episodes were recorded with home cage monitoring and were often observed immediately prior to their sudden death (Extended Data Video 1). Weakness, occasional pulling/paralysis of hindlimb, and uncoordinated walking appeared with increasing age, culminating in death. Weakness of the *Tmem65^-/-^* mice was also demonstrated by the lack of grip strength in an inverted grid test at P21 (Fig. 1h). Conversely, while the smaller hearts and lower heart rates were consistent with the growth delay^16^, no major cardiac structural or functional abnormalities were observed in the *Tmem65^-/-^* mice (Extended Data Figure 4). These data indicate an indispensable role for TMEM65 in mice and that the *Tmem65^-/-^* mouse model recapitulates the mitochondrial encephalomyopathy without cardiomyopathy phenotype observed clinically^1^.

## Loss of TMEM65 in neurons is lethal

Based on the high TMEM65 expression in the brain, seizures preceding death in TMEM65 KO mice, and the reported clinical phenotype^1^, we hypothesized that the primary functional impact of whole-body TMEM65 loss was on the brain. Indeed, magnetic resonance imaging (MRI) of P20-23 brains from Tmem65^+/+^ and *Tmem65^-/-^* littermates revealed lesions and a microcephaly phenotype in TMEM65 KO mice with the most apparent volume loss in the cerebral cortex and midbrain (Fig. 2a,b), consistent with a pathological loss of neurons^17^. Additionally, neuronal vacuolar degeneration accompanied by nuclear shrinkage and condensation was observed in the cingulate cortex (Fig. 2c) and brain stem (Extended Data Fig. 5) on H&E stained sagittal sections. Interestingly, neuronal vacuolar degeneration has been previously reported in Leigh syndrome, another mitochondrial encephalomyopathy often accompanied by seizures^18^, and thus may be related to the phenotypes associated with TMEM65 KO.

**Figure 2.**
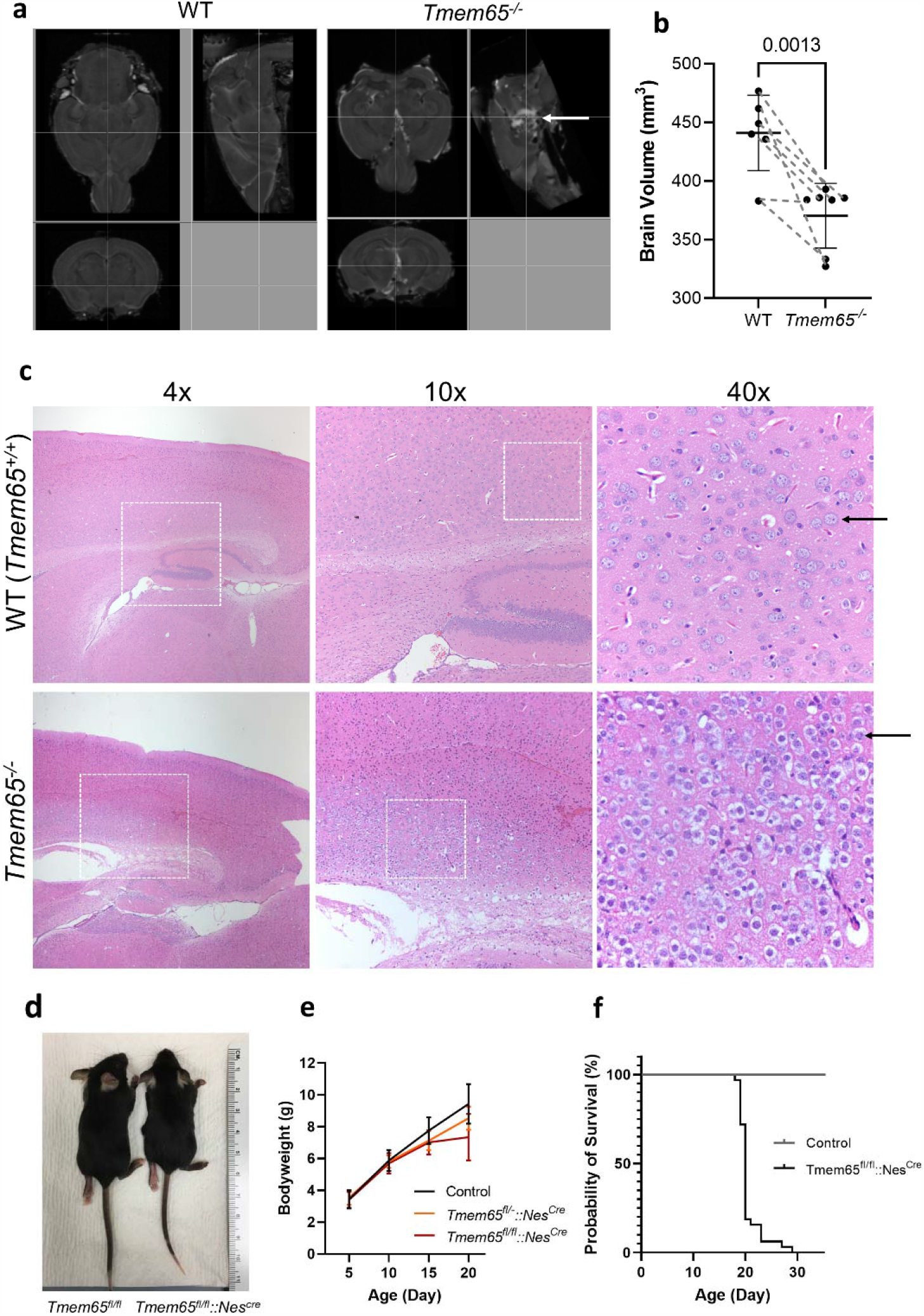
Loss of TMEM65 in brain causes neuronal vacuolar degeneration and sudden death. **a**, MRI scan of a pair of brains from P23 littermates show lesion and deformation of the *Tmem65^-/-^* brain. White arrow indicates lesion in midbrain. **b**, Mouse brain volumes measured with MRI scan. Dash line indicates littermate pairs. WT n = 6; *Tmem65^-/-^* n = 7. Individual values as well as mean ± SD are presented. **c**, H&E staining of brain sagittal sections show diffusive neuronal vacuolar degeneration in the cingulate cortex region. Black arrows indicate normal and degenerated neurons in brain sections. **d**, Representative image of *Tmem65^fl/fl^* control and *Tmem65^fl/fl^*::*Nes^Cre^* littermates at P20. **e**, Body weights of littermates measured at P5, P10, P15 and P20. Control (Tmem65^fl^ or *Tmem65^fl/fl^*) n = 25-36; Tmem65^fl^::*Nes^Cre^* n = 16-20; *Tmem65^fl/fl^*::*Nes^Cre^* n = 13-22. All data are presented as mean ± SD. **f**, Survival chart of controls and Tmem65^flox/flox^ ::*Nes^Cre^* pups.

To test whether loss of TMEM65 in the brain causes the sudden death observed in whole-body TMEM65 KO mice, TMEM65 was specifically ablated in the mouse nervous system by crossing the *Tmem65^fl/fl^* mouse with the Nestin-Cre mouse line^19^, in which Cre recombinase expression starts at embryonic day (E) 12.5 in the central and peripheral nervous system^20^. *Tmem65^fl/fl^::Nes^Cre^* mice recapitulated the epilepsy and sudden death in *Tmem65^-/-^* mice as all died between P18-P29 with an average life span of 20.5 ± 2.3 days (Fig. 2f) and also developed growth delays starting at P15 (Fig. 2d, 2e). The phenotypic similarity between the whole-body KO *Tmem65^-/-^* mice and nervous system-specific KO *Tmem65^fl/fl^::Nes^Cre^* mice confirms that TMEM65 is indispensable for normal brain structure and function.

## TMEM65 mediates adult-onset muscle loss

Gross motor defects and muscle weakness, as described above for the whole-body TMEM65 KO mice, are most commonly attributable to dysfunctional skeletal muscle, in addition to brain^21,22^. To assess whether loss of TMEM65 in skeletal muscle accounts for the gross motor defects in whole-body TMEM65 KO mice, we generated skeletal muscle-specific TMEM65 KO mice by crossing *Tmem65^fl/fl^* mice with Myf6-Cre mice where Cre expression is restricted to skeletal muscle starting in late gestation embryos^23-25^. The resultant *Tmem65^fl/fl^*::*Myf6^Cre^* mice lacking TMEM65 in skeletal muscle (Fig. 3a) are viable, fertile, and maintain normal bodyweight (Fig. 3b) and muscle fiber type distribution (Extended Data Figure 6a,c) at 2 months of age. Thus, loss of TMEM65 in skeletal muscle alone does not account for the early-onset motor dysfunction in whole-body TMEM65 KO mice.

**Figure 3.**
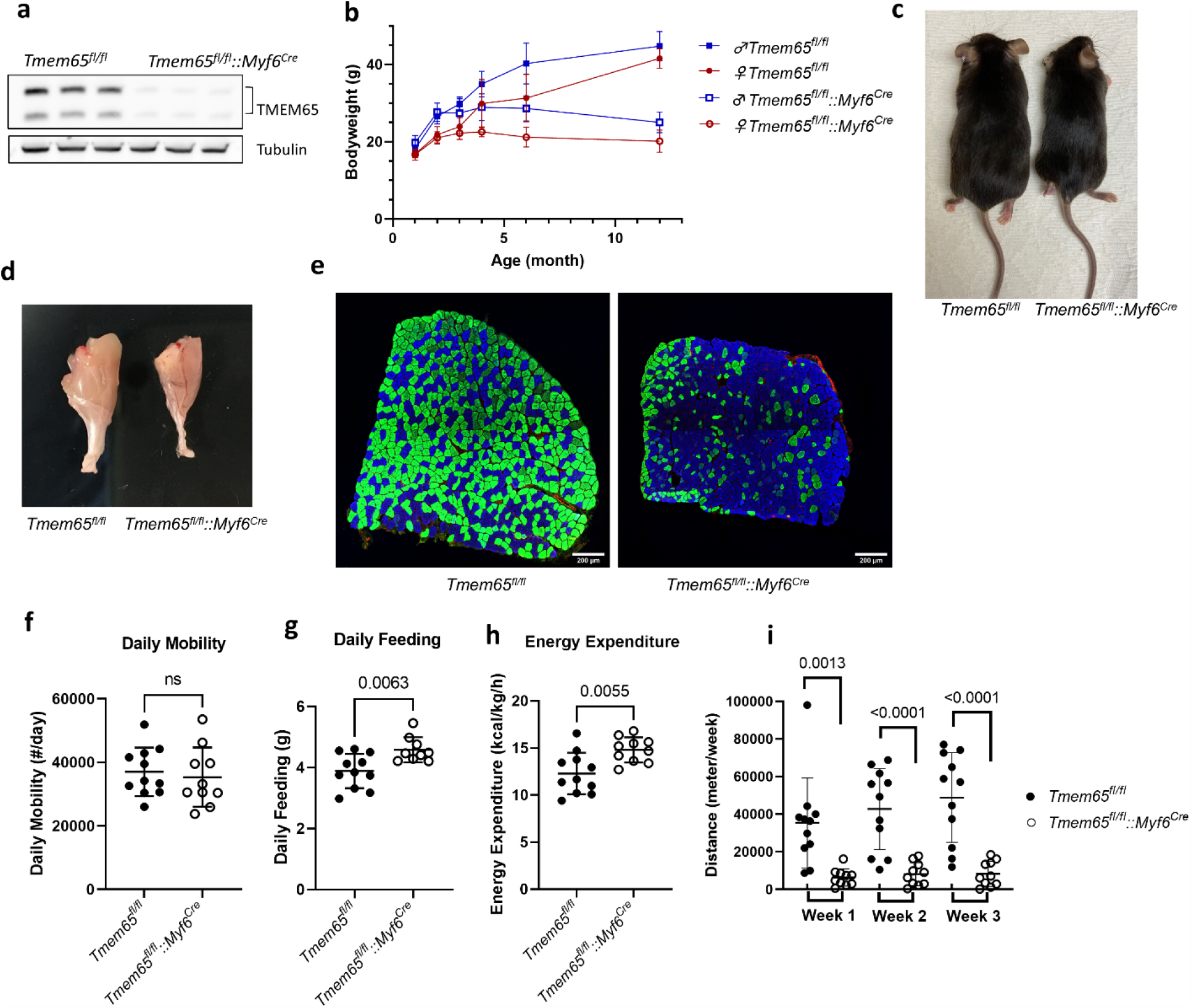
Loss of TMEM65 in skeletal muscle affects metabolism and leads to muscular atrophy and body weight loss. **a**, Western blots of soleus muscles from 2 months old *Tmem65^fl/fl^* and *Tmem65^fl/fl^*::*Myf6^Cre^* littermates show loss of TMEM65 in *Tmem65^fl/fl^*::*Myf6^Cre^* muscles. **b**, Bodyweights of male and female mice at different ages. Female *Tmem65^fl/fl^* n = 5-24; female *Tmem65^fl/fl^*::*Myf6^Cre^* n = 10-30; male *Tmem65^fl/fl^* n= 8-18; male *Tmem65^fl/fl^*::*Myf6^Cre^* n = 11-14. All data are presented as mean ± SD. **c**, Representative images of a pair of *Tmem65^fl/fl^* and *Tmem65^fl/fl^*::*Myf6^Cre^* littermates at 6-month-old. **d**, Gastrocnemius muscles from 6-month-old *Tmem65^fl/fl^* and *Tmem65^fl/fl^*::*Myf6^Cre^* littermates. **e**, Immunofluorescence of soleus muscles from *Tmem65^fl/fl^* and *Tmem65^fl/fl^*::*Myf6^Cre^* littermates at 6-month-old. Myosin isoforms were identified with specific antibodies to muscle type I (blue), IIa (green) and IIb (red). **f-h**, Metabolic parameters from 6-month-old control *Tmem65^fl/fl^* and *Tmem65^fl/fl^*::*Myf6^Cre^* littermates collected in metabolic cages (CLAMS) with simultaneous measurements. *Tmem65^fl/fl^* n = 11; *Tmem65^fl/fl^*::*Myf6^Cre^* n = 10. **i**, Voluntary wheel running tests of 6-month-old control *Tmem65^fl/fl^* and *Tmem65^fl/fl^*::*Myf6^Cre^* littermates collected in individual housing for 3 weeks. *Tmem65^fl/fl^* n = 11; *Tmem65^fl/fl^*::*Myf6^Cre^* n = 10. Individual value as well as mean ± SD are presented.

Despite the lack of overt phenotypes in 2 month old, muscle-specific TMEM65 KO mice, both male and female mice show growth delays by 3 months of age and regression after 4 months (Fig. 3b, 3c). By 6 months, *Tmem65^fl/fl^*::*Myf6^Cre^* mice have significantly lower body weight (Fig. 3b, 3c), muscle mass (Fig. 3d), and muscle fiber cross-sectional area (CSA, Extended Data Figure 6a,b) compared to littermate controls (*Tmem65^fl/fl^*). Weight and muscle loss, frequently accompanied by hunched body posture (kyphosis), continued through one year of age (Fig. 3b, Extended Data Figure 6a,b,d) at which point mice were euthanized due to their emaciated nature. Adult-onset muscle contractile type switching is also prevalent in *Tmem65^fl/fl^*::*Myf6^Cre^* mice as evident by the loss of muscle fibers expressing the fast-twitch myosin IIa isoform together with an increase in slow-twitch type I myosin expressing fibers at 6 and 10 months (Fig. 3e, Extended Data Figure 6a,c). This preferential loss of type IIa muscle fibers is consistent with the relatively greater expression of TMEM65 in this fiber type^26^ and demonstrates that persistent loss of TMEM65 in skeletal muscle leads to fast-to-slow fiber type switching and muscle atrophy.

To evaluate the functional impact of muscle-specific TMEM65 KO, we assessed whole body metabolism and exercise activity in 6 month old muscle-specific TMEM65 KO mice (*Tmem65^fl/fl^*::*Myf6^Cre^*) and littermate controls (*Tmem65^fl/fl^*). *Tmem65^fl/fl^*::*Myf6^Cre^* mice and controls were similarly active when housed in metabolic cages (Fig. 3f). However, daily feeding and energy expenditure were higher in *Tmem65^fl/fl^*::*Myf6^Cre^* mice compared to controls (Fig. 3g, 3h), indicating a reduced metabolic efficiency. Further, when given access to a voluntary running wheel for three weeks, *Tmem65^fl/fl^*::*Myf6^Cre^* mice ran less than 20% of their littermate controls each week (Fig. 3i). The reduction in metabolic efficiency and lack of exercise performance combined with the loss of both fat and lean mass (Extended Data Figure 6d) are all consistent with a metabolic uncoupling phenotype^27^ in mice lacking skeletal muscle TMEM65.

## TMEM65 regulates mitochondrial calcium efflux

TMEM65 was previously identified as a candidate mitochondrial calcium channel during the identification of the mitochondrial calcium uniporter (MCU)^28^. Indeed, the TMEM65 protein structure contains a glycine zipper motif^6^ which is known to facilitate formation of transmembrane channel proteins^29^, and kinetoplastid orthologs of TMEM65 contain a domain for a calcium binding EF-hand (Subfamily Protein Architecture Labeling Engine, SPARCLE^30^). Additionally, seizures^31^, gross motor defects^31^, apoptosis^31,32^, muscle atrophy^31^, and metabolic uncoupling phenotypes^32^, as described here for the TMEM65 KO mice, have all previously been linked to mitochondrial calcium handling. Indeed, mitochondrial calcium levels in isolated muscle fibers from 2 month old *Tmem65^fl/fl^*::*Myf6^Cre^* mice were higher than in littermate controls (Fig. 4a) and occurred without any changes in expression of known mitochondrial or cellular calcium handling proteins or metabolic pathways (Figs. 3a,4b, Extended Data Figure 7a-e). Thus, the increase in mitochondrial calcium level preceded the onset of the metabolic uncoupling phenotype and major protein expression changes that occur later in *Tmem65^fl/fl^*::*Myf6^Cre^* mice (Figs. 3,4b, Extended Data Figures 6,7). Mitochondrial calcium uptake rate was similar, if not slower, in isolated mitochondria from 2 month *Tmem65^fl/fl^*::*Myf6^Cre^* mice compared to controls (Fig. 4c,d) and mitochondrial calcium retention capacity was also unaffected (Fig. 4c,d). Additionally, isolated mitochondrial calcium efflux in the absence of sodium was similar between 2 month *Tmem65^fl/fl^*::*Myf6^Cre^* mice and controls (Fig. 4e,f). However, upon the addition of sodium, calcium efflux rate increased significantly in controls, but not in mitochondria from *Tmem65^fl/fl^*::*Myf6^Cre^* mice (Fig. 4e,f). Similarly, sodium-dependent mitochondrial calcium efflux is also reduced in brain mitochondria from 19 day old whole body TMEM65 KO mice despite no change in protein levels of the mitochondrial calcium import channel, MCU, or putative mitochondrial calcium exporters, NCLX, GHITM (aka TMBIM5/MICS1), and LETM1 (Extended Data Figure 8). These data indicate that TMEM65 regulates sodium-dependent mitochondrial calcium export both before and after the onset of the major TMEM65 KO-induced phenotypes in mouse tissues.

**Figure 4.**
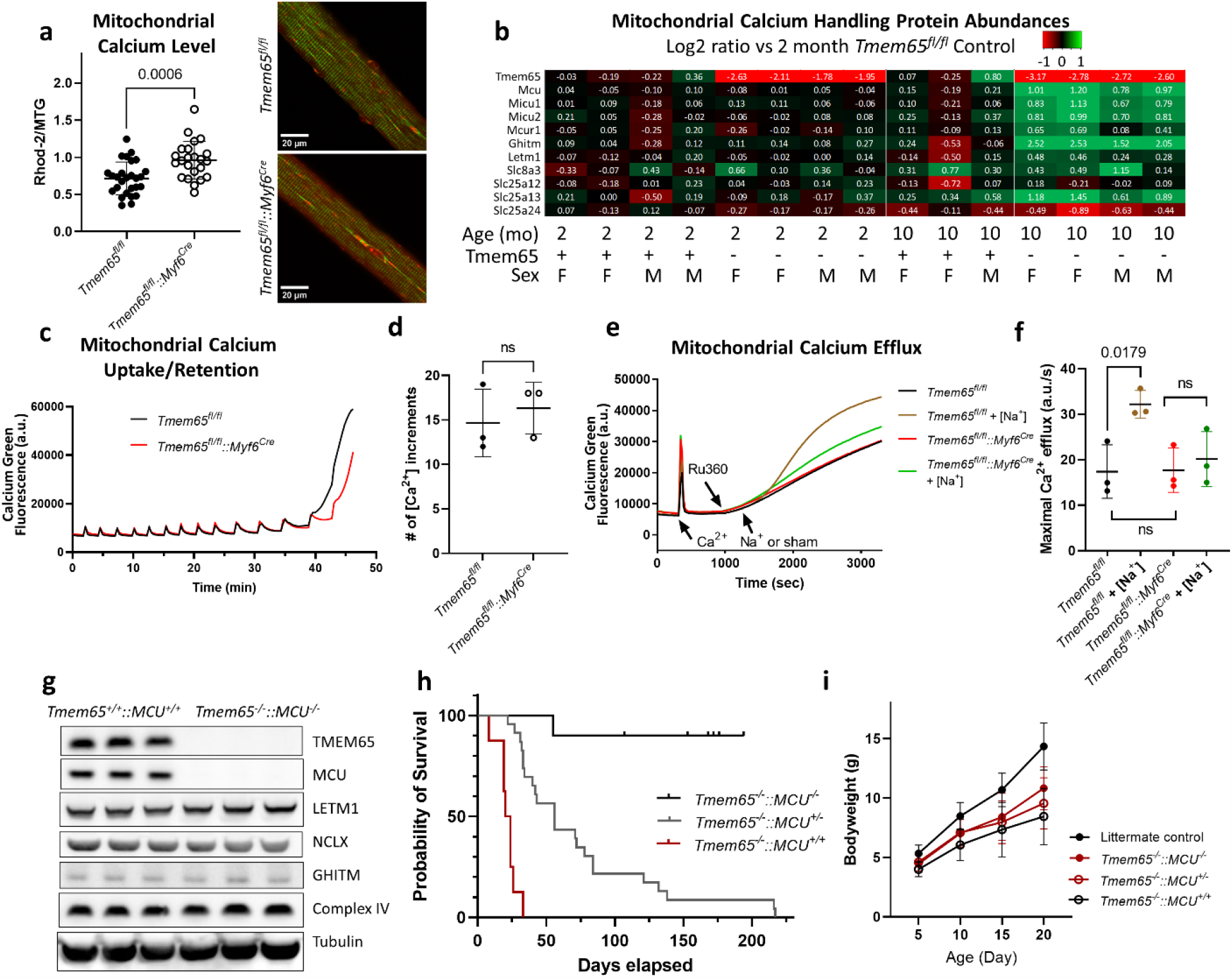
TMEM65 regulates mitochondrial calcium release in Na dependent manner. **a**, Relative mitochondrial matrix calcium level in FDB muscle from control *Tmem65^fl/fl^* (n = 27 cells from 3 mice) and *Tmem65^fl/fl^*::*Myf6^Cre^* (n = 23 cells from 3 mice) as measured by the fluorescence ratio of Rhod-2 to MitoTracker Green. Individual values as well as mean ± SD are presented. Representative images of *Tmem65^fl/fl^* and *Tmem65^fl/fl^*::*Myf6^Cre^* FDBs are shown on the right. **b**, Protein expression levels of mitochondrial calcium transport channels, carriers, and regulators detectable by mass spectrometry of soleus muscles from 2-month-old and 10-month-old *Tmem65^fl/fl^* or *Tmem65^fl/fl^*::*Myf6^Cre^* mice. **c**, Representative traces of extramitochondrial calcium with sequential Ca^2+^ additions (5µM) to isolated skeletal muscle mitochondria (100 µg) of *Tmem65^fl/fl^* and *Tmem65^fl/fl^*::*Myf6^Cre^* mice. **d**, Summary of the mitochondrial calcium retention capacity of isolated skeletal mitochondria of *Tmem65^fl/fl^* and *Tmem65^fl/fl^*::*Myf6^Cre^* mice. n = 3. **e**, Representative traces of mitochondrial calcium efflux assay. A bolus of Ca^2+^ (50 µM), MCU inhibitor, Ru360 (3 µM), and NaCl (20 mM) or sham were added sequentially to isolated skeletal muscle mitochondria (100 µg/well) incubated with Calcium Green. **f**, Summary of the maximal rates of mitochondrial Ca^2+^ efflux induced by Ru360 alone, or Ru360 and Na^+^ in *Tmem65^fl/fl^* control or *Tmem65^fl/fl^*::*Myf6^Cre^* mitochondria. n = 3 replicates per group. **g**, Western blot analysis of brain tissues from P20 WT (Tmem65^+/+^::MCU^+/+^) and *Tmem65^-/-^*::*MCU^-/-^* confirmed the loss of both TMEM65 and MCU in *Tmem65^-/-^*::*MCU^-/-^* mice. Expression level of other calcium handing proteins does not change. n = 3 replicates per group. **h**, Survival chart for *Tmem65^-/-^*::MCU^+/-^ (n = 24), *Tmem65^-/-^*::*MCU^-/-^* (n = 10) and *Tmem65^-/-^*::MCU^+/+^ (n = 8) mice. **i**, Bodyweights of control (n = 86-129), *Tmem65^-/-^*::MCU^+/+^ (n = 4-5) *Tmem65^-/-^*::MCU^+/-^ (n = 21-25) and *Tmem65^-/-^* ::*MCU^-/-^* (n = 5-6) mice before weaning. Data are represented as mean ± SD.

## Rebalancing calcium restores lifespan

We then hypothesized that rebalancing mitochondrial calcium levels by inhibiting mitochondrial calcium uptake would rescue the lethality of TMEM65 KO. Thus, we crossed *Tmem65^-/-^* mice with global MCU knockout mice^33^ (*MCU^-/-^*) which led to complete loss of TMEM65 and MCU without altering the abundances of LETM1, GHITM, NCLX, or mitochondrial complex IV in 20 day old brains of *Tmem65^-/-^* ::*MCU^-/-^* mice (Fig. 4g). Whole body TMEM65 KO again led to death around 20 days of age (Fig. 4h) indicating that the mixed genetic background (C57BL/6 and CD1) of the *Tmem65^-/-^*::*MCU^+/+^* mice did not alter the phenotype. However, heterozygous MCU KO (*Tmem65^-/-^*::*MCU^+/-^*) more than doubled the lifespan of TMEM65 KO mice while homozygous MCU KO (*Tmem65^-/-^*::*MCU^-/-^*) further prolonged lifespan (Fig. 4h) despite not fully rescuing the delayed growth associated with TMEM65 KO (Fig. 4i). These results demonstrate that the lethality associated with TMEM65 KO is mediated by mitochondrial calcium handling.

## Discussion

Tissue-specific variability in the expression of TMEM65 and other known mitochondrial calcium import/export proteins^34-38^ highlights the complexity of mitochondrial calcium handling within different parts of the body and points to the need for more comprehensive understanding of its regulatory mechanisms. The conditional TMEM65 KO mouse model developed here offers compelling evidence that loss of sodium-dependent mitochondrial calcium export in the brain leads to a lethal encephalopathy phenotype. This broadens the existing framework of knowledge beyond the accelerated progression of Alzheimer’s disease and cognitive decline associated with the neuronal-specific deletion of the putative mitochondrial sodium-dependent calcium exporter, NCLX^39,40^, and adds another dimension to the role of mitochondrial calcium in neuronal health. The different phenotypes following TMEM65 and NCLX deletion in neurons suggest an interesting avenue for future investigation, particularly in how or whether these two proteins may work together to mediate the transport of calcium out of mitochondria in support of neuronal function.

The adult-onset of the muscle atrophy and tissue remodeling phenotype upon loss of TMEM65 in skeletal muscle provides a unique window of time when the primary functions of TMEM65 can be probed without major influence from the pathological adaptations that occur with whole-body and neuronal-specific TMEM65 ablation. In this model, loss of TMEM65 results in greater accumulation of mitochondrial calcium and a lack of sodium-stimulated mitochondrial calcium efflux despite no alterations in the expression of known mitochondrial calcium handling machinery. Our data corroborate recent work showing that mitochondrial sodium/calcium exchange is activated when TMEM65 is overexpressed and blocked when TMEM65 is removed in vitro (accompanying DeStefani/Elrod papers). That rebalancing mitochondrial calcium dynamics by removing MCU is able to rescue the lethality associated with loss of TMEM65 in the brain offers further support for the importance of TMEM65 in mediating mitochondrial calcium export. These results link disruption in mitochondrial calcium homeostasis to severe mitochondrial disease and suggest inhibition of mitochondrial calcium influx^41^ may be a potential therapeutic target for defects associated with TMEM65 loss of function^1^ or other diseases characterized by mitochondrial calcium overload^39,40,42,43^.

## Methods

### Generation of transgenic mice

All mouse proced

ures were performed in accordance with institutional guidelines for animal studies and were approved by the National Heart, Lung, and Blood Institute Animal Care and Use Committee. To investigate the in vivo function of Tmem65 gene, we first attempted to generate a mouse line with total body Tmem65 knockout by co-microinjecting Cas9 mRNA with two single guide RNAs (sgRNAs) targeting Exon 1 and Exon 4 (the italic and underlined nucleotides are PAM):

Tmem-Ex1: gccgtgttcagcgcctccat<u>*ggg*</u>

Tmem-Ex4: gcgatacctttcgtagggtt<u>*tgg*</u>

into zygotes collected from C57BL/6N mice, but all of the 19 offspring died between 2- and 4-week of age. Since hybrid mice show hybrid vigor and sometimes can help overcoming lethality, we next injected the same CRISPR reagents into zygotes collected from B6CBA/F1 hybrid mice (JAX #100011), but again all of the 26 offspring died between 2 and 4 weeks. Some of these offspring were genotyped by PCR either shortly before or shortly after their death, and all of them turned out to be homozygous mutants, indicating that these pair of sgRNAs are extremely potent at making deletions at their target sites. In attempting to obtain heterozygous mutant mice by reducing CRISPR cutting potency, we then microinjected Cas9 mRNA with each of these two sgRNAs alone. All of the 18 offspring injected with Tmem-Ex4 sgRNA died, and all but one of the 24 pups injected with Tmem-Ex1 sgRNA died before they are one month old. The single surviving mouse appeared normal and bred well. However, PCR and sequencing analyses of this founder and its offspring revealed that this line had a 3bp deletion in Exon 1, which led to the in frame deletion of E58 from the TMEM65 protein. Therefore, it is a point mutation mouse line rather than a Tmem65 knockout.

The TMEM65 conditional KO mouse line was generated by sequentially inserting loxP sites into Intron 3 and Intron 4, so that Exon4 can be excised by exposing to Cre recombinase, resulting to translation frame shift. Briefly, the Intron 3 loxP was inserted by co-microinjecting Cas9 mRNA with the Tmem-In3 sgRNA:

gtagttttgtgatagtcctc<u>agg</u>

and the Tmem-In3 donor oligos:

gaatgtaggcatttttaaaaattattattacaaatacagagattagtagttttgtgatagtataacttcgtatagcatacattatacgaagttatggatcccctcaggcacaagagagaaatagactggcatgtgtcaggttaaatgggggcatcccgtg

into zygotes collected from C57BL6/N mice. The Intron 4 loxP was inserted by co-microinjecting Cas9 mRNA with the Tmem-In4 sgRNA:

cattctagaagtacagctag<u>agg</u>

and the Tmem-In4 donor oligos:

tcacgttgcaacatcttctgccttcgttgtgttgtgtagtattatgaaaagttccctctagataacttcgtatagcatacattatacgaagttataagcttctgtacttctagaatgctttatggattcaactattttggatattagcagtaaatagactg

into zygotes collected from mice that already contain the Intron 3 loxP site. Mice containing both loxP sites on the same allele were expanded and use for conditional knockout experiments.

To generate whole-body TMEM65 KO mice, *Tmem65^fl/fl^* mice were crossed with CMV-Cre transgenic mice (available from the Jackson Laboratory, stock no. 006054). To generate neuronal specific TMEM65 KO mice, *Tmem65^fl/fl^* mice were crossed with Nestin-Cre transgenic mice (the Jackson Laboratory, stock no. 003771). To generate skeletal muscle-specific TMEM65 KO mice, *Tmem65^fl/fl^* mice were crossed with Myf6-Cre mice (the Jackson Laboratory, stock no. 010528).

The TMEM65-V5 mouse line was generated by adding a V5 tag sequence in-frame to the c-terminus of the Tmem65 gene, immediately before the stop codon. Briefly, Cas9 mRNA was co-microinjected with the Tmem-V5 sgRNA:

atcttcttataggtgttcta<u>agg</u>

and the Tmem-V5 donor oligos:

ggaatgttcccgttgattttctttggaggaagtgaagaggatgagaaactggaaacaacaaatggcaagcccatccccaaccccctgctgggcctggacagcacctaatcacgtttcaacacctataagaagatgtaaactaatgtacctcatcattaactatactgtccccacagttagc

into zygotes collected from the Tmem65 conditional knockout mice. Mice containing both an intact V5 tag and correctly floxed Exon 4 were bred and used for experiments.

### Mouse line genotyping

Genomic DNA was extracted from mouse tail snips and standard PCR was performed for genotyping. The following primers were used for genotyping floxed mice:

5’ floxed site:

Forward: aaggactgcacagagatgtc

Reverse: acttgacaggagttatgctg

3’ floxed site:

Forward: cattctagactaccttaggtg

Reverse: gttagactctgtgaagctcac

Cre mice were genotyped according to the protocols from the Jackson Laboratory.

### Western Blot analysis

Mouse tissues were harvested and lysed mechanically with homogenizer in 1x RIPA lysis buffer (20-188, Sigma-Aldrich) containing 1x proteinase inhibitor cocktail (P8340, Sigma-Aldrich). Tissue lysates were incubated on ice for 10 minutes, and clarified by centrifugation at 10,000 x g for 10 minutes at 4°C to pellet the tissue debris. Supernatants containing equal amounts of soluble proteins were mixed with 2x SDS protein sample buffer containing 100mM DTT (Sigma-Aldrich), denatured by incubating on a heat block at 98 C for 5 minutes, and were loaded on NuPAGE 10% Bis-Tris protein gels (Life Technologies). The gel was running at 150 V for about one hour to separate the proteins. Proteins were then transferred from Bis-Tris gel onto PVDF membrane via XCell II blot module (Invitrogen) for immunoblotting. The PVDF membrane blots were blocked with blocking buffer (Li-Cor P/N927-40000) for one hour, incubated in blocking buffer containing 0.1% Tween and primary antibodies for 3 hours at room temperature or overnight at 4°C, washed 3 times with 1x PBS containing 0.1% Tween (PBST), incubated with blocking buffer containing 0.1% Tween and IRDye secondary antibodies (Li-Cor), washed 3 times with PBST, and imaged with Azure c600 imaging system (Azure Biosystems). Chameleon duo pre-stained protein ladder (Li-Cor 928-60000) was used as a reference of molecular weights for detected proteins. The primary antibodies used were anti-TMEM65 (HPA025020, Sigma-Aldrich, 1:400; PA5-112762, Invitrogen, 1:1K), V5 (R960-25, Life Technologies, 1:2K), α-tubulin (P/N 926-42212, Li-Cor, 1:2K), MCU (HPA016480, Sigma-Aldrich, 1:1K), LETM1 (16024-1-AP, Proteintech, 1:1K), NCLX (21430-1-AP, Proteintech, 1:1K; ab136975, Abcam, 1:1K), GHITM (16296-1-AP, 1:500), Complex IV subunit IV (A21348, Invitrogen, 1:1K).

### Tissue harvesting and fixation for H&E staining

Under anesthesia, P20 or older mice were perfused with 1xPBS solution, followed by perfusion of 1xPBS containing 4% paraformaldehyde (PFA). Mouse Brain tissue was dissected out and fixed in 1xPBS containing 4% PFA at 4 °C overnight. The fixed brain tissue was washed with 1xPBS, and stored in 70% ethanol for paraffin embedding. Paraffin embedding, tissue sectioning, and H&E staining were performed by Histoserv Inc (Germantown, MD).

### Confocal microscopy

Mouse was anesthetized with 5% Isoflurane through gas anesthesia system, and was perfused through heart with 1xPBS and then 4% paraformaldehyde in 1xPBS. Mouse tissue was harvested and further fixed for 4 hours. Fixed tissue was washed with 1x PBS, and transferred into 30% sucrose in 1xPBS solution overnight till tissue sank to the bottom of the solution. The tissue was trimmed and embedded in optimal cutting temperature (O.C.T.) compound (Sakura). For skeletal muscles, tissue were immediately embedded in O.C.T. after harvesting. O.C.T. embedded tissues were then quickly frozen in isopentane prechilled in dry ice. 10 µm sections were cut in cryostat at -20 °C. Frozen tissue sections were blocked with blocking buffer (1xPBS containing 5% goat serum and 1% bovine serum albumin) for one hour, and then incubated in blocking buffer containing 0.05% Tween and primary antibodies overnight. Sections were washed 3 times with PBS containing 0.05% Tween, and then incubated in Alexa Fluor secondary antibodies (Invitrogen) for one hour. Coverslips were mounted on the sections with coverslips with Prolong gold antifade mountant (Life Technologies) after 3 washes of 1xPBS containing 0.05% Tween. Confocal images were collected with a Zeiss LSM 780 confocal microscope system. Primary antibodies used were to ATP5A (ab14748, Abcam, 1:200), V5 (13202S, Cell Signal Technology, 1:100; or R960-25, Life technologies, 1:200), TMEM65 (HPA025020, Sigma-Aldrich, 1:50), Desmin (D1033, Sigma-Aldrich, 1:100), Connexin 43 (ab11370, Abcam, 1:400), Myosin heavy chain type I (BA-D5, DSHB, 1:60), Myosin heavy chain type IIa (SC-71, DSHB, 1:100), Myosin heavy chain type IIb (BF-F3, DSHB, 1:80), Laminin (L9393, Sigma-Aldrich, 1:200).

### Inverted grip suspension test

7 wild type (WT) and 6 KO mice of P21 were subject to inverted grip strength tests. Mouse was gently placed on top of a wire grid to allow its paws to attach the grid with its torsi horizontal. The grip was gently inverted 180 degrees so the mouse was suspended above a padded surface. Latency to fall was record and averaged for three trials, separated by a rest period. The maximum trial length was set to 180 seconds.

### Home cage monitoring

Paired breeding mice with new born liters containing at least 1 WT and 1 KO pups were moved into PhenoTyper home cages (Noldus, Wageningen, Netherlands) equipped with camera mounted in the cage lids for 24 hour video monitoring to record episodes of epilepsy and any other behavior changes in the pups.

### Indirect calorimetry

The Oxymax-CLAMS setup (Columbus Instruments) was used to assess mouse metabolism. Mice were singly housed in the CLAMS chambers, received food and water ad libitum throughout the experiment, and were checked on at least twice a day throughout the experiment. Oxygen consumption, carbon dioxide production, food intake, and beam breaks (locomotion) were monitored during the experiment. The resulting data was analyzed using CalR.

### Body composition

Whole body lean and fat mass was measured using the EchoMRI™ body composition analyzers for live mice.

### Magnetic resonance imaging (MRI)

A total of thirteen 20, 21, and 23 days old littermates of WT and KO pups (7 KOs and 6 WTs) were anesthetized with 5% Isoflurane through gas anesthesia system and perfused with 1xPBS, and then 1xPBS containing 4% PFA and Magnevist (1:500, Bayer HealthCare Pharmaceuticals) to enhance the brain with MR contrast. The mouse head was dissected out and the skin was removed to obtain the brain within the skull. The samples were then stored in 1xPBS containing 4% PFA in 4°C refrigerator. The brain remained inside the cranium for scanning to preserve the brain anatomy. Before the scan, samples were washed and equilibrated in 1xPBS for at least 24 hours.

MRI acquisition was performed ex vivo on a 14T Bruker (Billerica, MA, USA) microimaging system with a 10mm linear radio frequency coil. A T2-weighted image was acquired using a 3D multislice multiecho (MSME) pulse sequence with 100 µm isotropic resolution and a matrix size of 160 × 84 × 80, and with TE/TR = 30/3000, nex = 1. To examine brain volume, masks for the brain parenchyma were created that excluded skull and other non-brain material using the automatic segmentation snake tool in ITK-SNAP^44^. The volumes of these parenchymal brain masks were measured using ITK-SNAP and the values were plotted for each specimen.

### Echocardiography and electrocardiography

Mouse heart echocardiography and electrocardiography (ECG) were performed at the NHLBI Phenotyping Core. Mice were lightly anesthetized with isoflurane and placed supine over a heated platform with ECG leads and a rectal temperature probe. The Vevo2100 ultrasound system (VisualSonics, Toronto, Canada) with a 30 MHz ultrasound probe (VisualSonics, MS-400 transducer) was used to acquire heart images. Measurements were made from standard 2D and M mode images from the parasternal long axis and mid-papillary short axis views of the left ventricle.

### Dobutamine stress echocardiography

Mice were lightly anesthetized with isoflurane during examination and placed supine on a heated platform with ECG leads and a rectal temperature probe. Heart images were acquired using the Vevo2100 ultrasound system (VisualSonics) with a 30-MHz ultrasound probe (VisualSonics, MS-400 transducer). After a baseline scan, mice were given a constant rate infusion of dobutamine (0.625 mg/ml in normal saline containing 5% dextrose) via the tail vein using an infusion syringe pump (Harvard Apparatus). The first infusion was at the low dose of 10 μg/kg/min. After the heart rate reached a steady state as determined by the ECG, infusion rate was increased to 40 μg/kg/min, and the scans were repeated. Echocardiograph scans were collected after the low and high dose infusions. Post imaging analysis was conducted using Vevo2100 analysis software to measure %EF, %FS, and cardiac chamber dimensions from 2-D and M-mode images.

### Mass spectrometry

Muscle tissue preparation and mass spectrometry was performed as previously described^45^. Briefly, frozen soleus muscles were individually transferred to 130 µl of urea based lysis buffer, 6 M Urea, 2 M Thiourea, 50 mM Triethylammonium bicarbonate (TEAB). Tissues were homogenized with ceramic beads using 3 cycles of 40s at 5500 rpm and 41C (Precellys® Cryolys Evolution, Bertin Technologies). Tissue lysates were further clarified by microcentrifuge spin columns (QIAshredder, Qiagen) to obtain clear protein lysate centrifuged at 10,000g for 2min at 41C. The extracted protein supernatants were transferred to 1.5 ml microtubes for further processing. Bradford assay (Thermo Fisher Scientific) was used to estimate the protein concentration of lysate, 100 µg of each lysate was reduced, alkylated, delipidated, digested with trypsin and individually labeled with Tandem Mass Tag (TMT) 16plex labeling reagent kit as per manufacturer’s instructions (Thermo Fisher Scientific).

High pH reversed-phase liquid chromatography was performed on an offline Agilent 1200 series HPLC. The quenched TMT labeled peptides were all combined, concentrated and desalted using HLB Oasis 1cc, Waters) column per manufacturer’s instructions. The dried desalted peptides were resuspended in 0.1 ml 10 mM triethyl ammonium bicarbonate with 2% (v/v) acetonitrile. Peptides were loaded onto an Xbridge C18 HPLC column (Waters; 2.1mm inner diameter × 100 mm, 5μm particle size), and profiled with a linear gradient of 5–35 % buffer B (90% acetonitrile, 10 mM triethyl ammonium bicarbonate) over 60 min, at a flowrate of 0.25 ml/min. The chromatographic performance was monitored by sampling the eluate with a diode array detector (1200 series HPLC, Agilent) scanning between wavelengths of 200 and 400 nm. Fractions were collected at 1min intervals followed by fraction concatenation. Twelve concatenated fractions were dried and resuspended in 0.01% formic acid, 2% acetonitrile. Approximately 500 ng of peptide mixture was loaded per liquid chromatography-mass spectrometry run.

All fractions were analyzed on an Ultimate 3000-nLC coupled to an Orbitrap Fusion Lumos Tribrid instrument (Thermo Fisher Scientific) equipped with a nanoelectrospray source. Peptides were separated on an EASY-Spray C18 column (75 μm × 50cm inner diameter, 2 μm particle size and 100 Å pore size, Thermo Fisher Scientific). Peptide fractions were placed in an autosampler and separation was achieved by 120 min gradient from 4-40% buffer B (100% ACN and 0.1% formic acid) at a flow rate of 300 nL/min. An electrospray voltage of 1.9 kV was applied to the eluent via the EASY-Spray column electrode. The Lumos was operated in positive ion data-dependent mode, using Synchronous Precursor Selection (SPS-MS3). Full scan MS1 was performed in the Orbitrap with a precursor selection range of 380–1,400 m/z at nominal resolution of 1.2 × 105. The AGC target and maximum accumulation time settings were set to 4 × 105 and 50 ms, respectively. MS2 was triggered by selecting the most intense precursor ions above an intensity threshold of 5 × 103 for collision induced dissociation (CID)-MS2 fragmentation with an AGC target and maximum accumulation time settings of 2 × 104 and 75 ms, respectively. Mass filtering was performed by the quadrupole with 0.7 m/z transmission window, followed by CID fragmentation in the linear ion trap with 35% normalized collision energy in rapid scan mode and parallelizable time option was selected. SPS was applied to co-select 10 fragment ions for HCD-MS3 analysis. SPS ions were all selected within the 350–1,300 m/z range and were set to preclude selection of the precursor ion and TMTC ion series. The AGC target and maximum accumulation time were set to 1 × 105 and 150 ms (respectively) and parallelizable time option was selected. Co-selected precursors for SPS-MS3 underwent HCD fragmentation with 55% normalized collision energy and were analyzed in the Orbitrap with nominal resolution of 5 × 104. The number of SPS-MS3 spectra acquired between full scans was restricted to a duty cycle of 3 s.

Raw data files were processed using Proteome Discoverer (v2.4, Thermo Fisher Scientific), with Sequest HT (Thermo Fisher Scientific) search node. The data files were searched against UniProtKB/Swiss-Prot mus musculus (17,030 reviewed, release 2020_10) protein sequence database, with carbamidomethylation of cysteine, TMT-Pro (+304.207) modification of lysines and peptide N-terminus set as static modifications; oxidation of methionine as dynamic. For SPS-MS3, the precursor and fragment ion tolerances of 10 ppm and 0.5 Da were applied, respectively. Up to two-missed tryptic cleavages were permitted. Percolator algorithm (v.3.02.1, University of Washington) was used to calculate the false discovery rate (FDR) of peptide spectrum matches (PSM), set to a q-value <0.05. TMT 16-plex quantification was also performed by Proteome Discoverer by calculating the sum of centroided ions within 20 ppm window around the expected m/z for each of the 15 TMT reporter ions. Spectra with at least 60% of SPS masses matching to the identified peptide are considered as quantifiable PSMs. Quantification was performed at the MS3 level where the median of all quantifiable PSMs for each protein group was used for protein ratios. The mass spectrometry proteomics data have been deposited to the ProteomeXchange Consortium (dataset identifier PXD035605) via MassIVE (UCSD, San Diego, CA, USA) a member of the consortium (dataset identifier MSV000089991).

### Muscle fiber isolation and calcium imaging

The FDB muscle was dissected from the mouse foot and digested in collagenase (Sigma) at 37°C for 2-3 hours. After digestion, individual fibers were released by gentle trituration in MEM culture medium, and plated on Matrigel-coated dishes. After 1 hour incubation with 4µM Rhod-2-AM and 100 nM MitoTracker Green (Thermo Fisher Scientific) at 37°C, the buffer was gently exchanged with MEM culture medium. Fibers were imaged 20 minutes after. Rhod-2 and MitoTracker Green fluorescence were simultaneously measured on a Zeiss LSM 780 confocal microscope.

### Isolation of mitochondria from tissues

For isolation of skeletal muscle mitochondria, hind limb muscles were harvested from 2-month-old mice, and transferred into ~10 ml isolation medium (150 mM sucrose, 75 mM KCl, 50 mM Tris-HCl, 1 mM KH2PO4, 5 mM MgCl2, 1 mM EGTA, pH 7.4) on ice. Muscles were minced into small pieces in isolation medium, and 0.5mg Subtilisin A (1mg/ml in isolation buffer, Sigma-Aldrich Cat# P3910) were added. Before homogenization, 5 ml of isolation medium containing with 0.2% BSA were added. The minced muscle tissues were then homogenized for 15 seconds at 40% of full power with Polytron (IKA Works, INC. Model: Ultra Turrax T25). Homogenate was centrifuged at 700g for 10 minutes at 4 °C. After first spin, supernatant was poured into a clean tube chilled on ice and then centrifugated at 10,000 g for 10 minutes at 4 °C. The resulting pellet (mitochondria) was resuspended in Buffer B (225mM Mannitol, 75mM Sucrose, 5mM MOPS, 2mM Taurine, 1mM EGTA, pH 7.25), 0.2% bovine serum albumin (BSA). The enrichment of mitochondria were wash and centrifuged at 11,000g for 3 min 4 °C. The final mitochondrial pellet was resuspended in 1 ml of Buffer B without albumin to measure Mitochondrial protein concentration by Bradford method.

For isolation of brain mitochondria, mouse brain were minced and then homogenized in ice-cold Buffer B containing 0.1% BSA. Homogenate was centrifuged at 700g for 10 minutes at 4 °C to pellet unlysed cells, and the supernatant was centrifugated at 10,000g for 10 minutes to pellet crude mitochondria. The crude mitochondria were resuspended in 3.5 mL of 15% Percoll solution in Buffer B, which was layered on top of 3.7 mL 24% and 1 mL 40% Percoll solutions in Buffer B. The solutions were then centrifugated at 30,000g for 40 minutes. Non-synaptosomal mitochondria were collected from a band between the 24% and 40% Percoll layers. Mitochondria were then resuspended in Buffer B and centrifuged at 10,000g for 10 minutes. The final mitochondrial pellet was resuspended in 1 ml of Buffer B without albumin to measure Mitochondrial protein concentration by Bradford method.

### Measurement of Ca^2+^ efflux from isolated mitochondria

Extramitochondrial Ca^2+^ concentration was monitored using the low-affinity fluorescent Ca^2+^ indicator Calcium Green 5N as previously described^33^. 100ug mitochondria were suspended in a respiration buffer (120mM KCl, 10mM Tris, 5mM MOPS, 5mM K_2_HPO_4_, 10mM L-glutamic acid, 5mM L-Malic acid, pH=7.4) in a 96 well, black/clear, tissue culture treated plate (Falcon). For a standard experiment, mitochondria were loaded with a bolus of Ca^2+^ (50µM), followed by addition of the MCU inhibitor Ru360 (3µM) to block mitochondrial Ca^2+^ uptake, thereby enabling observation of mitochondrial Ca^2+^ efflux in isolation. To probe Na^+^ dependent Ca^2+^ release from mitochondria, 20mM of NaCl was added to mitochondria in the presence of Ru360. Calcium Green 5N signal was recorded with a CLARIOstar Plus plate reader (BMG Labtech).

## Statistics

Results are expressed as mean ± SD. Statistical analyses were performed using GraphPad Prism 9 software. P value was reported on each test. A P value of less than 0.05 was considered statistically significant.

## Study approval

Animal studies were approved by the National Heart, Lung, and Blood Institute Animal Care and Use Committee.

## Author contributions

YZ, JS, EM, and BG designed research studies. YZ, BG, LR, CL, AN, AA, DS, JM, and RC conducted experiments. YZ, LR, AA, BG, and DS analyzed data. YZ and BG wrote the manuscript. All authors edited and approved final manuscript.

## Acknowledgements

The authors thank Robert Balaban and Vamsi Mootha for help with initial screens which identified Tmem65 as a gene of interest.

## References

1 Nazli, A. et al. A mutation in the TMEM65 gene results in mitochondrial myopathy with severe neurological manifestations. Eur J Hum Genet 25, 744–751 (2017). 10.1038/ejhg.2017.20

2 Pagliarini, D. J. et al. A mitochondrial protein compendium elucidates complex I disease biology. Cell 134, 112–123 (2008). 10.1016/j.cell.2008.06.016

3 Nishimura, N., Gotoh, T., Oike, Y. & Yano, M. TMEM65 is a mitochondrial inner-membrane protein. PeerJ 2, e349 (2014). 10.7717/peerj.349

4 Foulds, C. E. et al. Research resource: expression profiling reveals unexpected targets and functions of the human steroid receptor RNA activator (SRA) gene. Mol Endocrinol 24, 1090–1105 (2010). 10.1210/me.2009-0427

5 Sasarman, F. et al. LRPPRC and SLIRP interact in a ribonucleoprotein complex that regulates posttranscriptional gene expression in mitochondria. Molecular biology of the cell 21, 1315–1323 (2010).

6 Sharma, P. et al. Evolutionarily conserved intercalated disc protein Tmem65 regulates cardiac conduction and connexin 43 function. Nat Commun 6, 8391 (2015). 10.1038/ncomms9391

7 Teng, A. C. T. et al. Tmem65 is critical for the structure and function of the intercalated discs in mouse hearts. Nat Commun 13, 6166 (2022). 10.1038/s41467-022-33303-y

8 Suzuki-Hatano, S. et al. AAV9-TAZ gene replacement ameliorates cardiac TMT proteomic profiles in a mouse model of Barth syndrome. Molecular Therapy-Methods & Clinical Development 13, 167–179 (2019).

9 Regan, J. A. et al. Phenome-Wide Association Study of Severe COVID-19 Genetic Risk Variants. Journal of the American Heart Association 11, e024004 (2022).

10 Zhang, B. et al. The chromatin remodeler CHD6 promotes colorectal cancer development by regulating TMEM65-mediated mitochondrial dynamics via EGF and Wnt signaling. Cell Discovery 8, 130 (2022).

11 Shimizu, H. & Nakayama, K. I. A 23 gene–based molecular prognostic score precisely predicts overall survival of breast cancer patients. EBioMedicine 46, 150–159 (2019).

12 Wagner, K. W. et al. KDM2A promotes lung tumorigenesis by epigenetically enhancing ERK1/2 signaling. The Journal of clinical investigation 123, 5231–5246 (2013).

13 Song, X. et al. Pan-Cancer Analysis of Prognostic and Immune Infiltrates for the TMEM65, Especially for the Breast Cancer. Evidence-Based Complementary and Alternative Medicine 2023 (2023).

14 Glancy, B. et al. Mitochondrial reticulum for cellular energy distribution in muscle. Nature 523, 617–620 (2015). 10.1038/nature14614

15 Schwenk, F., Baron, U. & Rajewsky, K. A cre-transgenic mouse strain for the ubiquitous deletion of loxP-flanked gene segments including deletion in germ cells. Nucleic Acids Res 23, 5080–5081 (1995). 10.1093/nar/23.24.5080

16 Sato, S. Quantitative evaluation of ontogenetic change in heart rate and its autonomic regulation in newborn mice with the use of a noninvasive piezoelectric sensor. Am J Physiol Heart Circ Physiol 294, H1708–1715 (2008). 10.1152/ajpheart.01122.2007

17 Burte, F., Carelli, V., Chinnery, P. F. & Yu-Wai-Man, P. Disturbed mitochondrial dynamics and neurodegenerative disorders. Nat Rev Neurol 11, 11–24 (2015). 10.1038/nrneurol.2014.228

18 Lake, N. J., Bird, M. J., Isohanni, P. & Paetau, A. Leigh syndrome: neuropathology and pathogenesis. J Neuropathol Exp Neurol 74, 482–492 (2015). 10.1097/NEN.0000000000000195

19 Liang, H., Hippenmeyer, S. & Ghashghaei, H. T. A Nestin-cre transgenic mouse is insufficient for recombination in early embryonic neural progenitors. Biol Open 1, 1200–1203 (2012). 10.1242/bio.20122287

20 Tronche, F. et al. Disruption of the glucocorticoid receptor gene in the nervous system results in reduced anxiety. Nat Genet 23, 99–103 (1999). 10.1038/12703

21 Lucas, B. R. et al. Interventions to improve gross motor performance in children with neurodevelopmental disorders: a meta-analysis. BMC pediatrics 16, 1–16 (2016).

22 Emery, A. E. The muscular dystrophies. The Lancet 359, 687–695 (2002).

23 Sambasivan, R. et al. Embryonic founders of adult muscle stem cells are primed by the determination gene Mrf4. Dev Biol 381, 241–255 (2013). 10.1016/j.ydbio.2013.04.018

24 Southard, S. et al. A series of Cre-ER(T2) drivers for manipulation of the skeletal muscle lineage. Genesis 52, 759–770 (2014). 10.1002/dvg.22792

25 Keller, C. et al. Alveolar rhabdomyosarcomas in conditional Pax3:Fkhr mice: cooperativity of Ink4a/ARF and Trp53 loss of function. Genes Dev 18, 2614–2626 (2004). 10.1101/gad.1244004

26 Kallabis, S. et al. High-throughput proteomics fiber typing (ProFiT) for comprehensive characterization of single skeletal muscle fibers. Skelet Muscle 10, 7 (2020). 10.1186/s13395-020-00226-5

27 Clapham, J. C. et al. Mice overexpressing human uncoupling protein-3 in skeletal muscle are hyperphagic and lean. Nature 406, 415–418 (2000).

28 De Stefani, D., Raffaello, A., Teardo, E., Szabo, I. & Rizzuto, R. A forty-kilodalton protein of the inner membrane is the mitochondrial calcium uniporter. Nature 476, 336–340 (2011). 10.1038/nature10230

29 Kim, S. et al. Transmembrane glycine zippers: physiological and pathological roles in membrane proteins. Proc Natl Acad Sci U S A 102, 14278–14283 (2005). 10.1073/pnas.0501234102

30 Lu, S. et al. CDD/SPARCLE: the conserved domain database in 2020. Nucleic acids research 48, D265–D268 (2020).

31 Garbincius, J. F. & Elrod, J. W. Mitochondrial calcium exchange in physiology and disease. Physiological Reviews 102, 893–992 (2022).

32 Brookes, P. S., Yoon, Y., Robotham, J. L., Anders, M. W. & Sheu, S. S. Calcium, ATP, and ROS: a mitochondrial love-hate triangle. Am J Physiol Cell Physiol 287, C817–833 (2004). 10.1152/ajpcell.00139.2004

33 Pan, X. et al. The physiological role of mitochondrial calcium revealed by mice lacking the mitochondrial calcium uniporter. Nat Cell Biol 15, 1464–1472 (2013). https://doi.org:ncb2868

34 Palty, R. et al. NCLX is an essential component of mitochondrial Na+/Ca2+ exchange. Proceedings of the National Academy of Sciences 107, 436–441 (2010).

35 Palty, R. et al. Lithium-calcium exchange is mediated by a distinct potassium-independent sodium-calcium exchanger. Journal of Biological Chemistry 279, 25234–25240 (2004).

36 Paillard, M. et al. Tissue-specific mitochondrial decoding of cytoplasmic Ca2+ signals is controlled by the stoichiometry of MICU1/2 and MCU. Cell reports 18, 2291–2300 (2017).

37 Patron, M., Granatiero, V., Espino, J., Rizzuto, R. & De Stefani, D. MICU3 is a tissue-specific enhancer of mitochondrial calcium uptake. Cell Death & Differentiation 26, 179–195 (2019).

38 Fieni, F., Bae Lee, S., Jan, Y. N. & Kirichok, Y. Activity of the mitochondrial calcium uniporter varies greatly between tissues. Nature communications 3, 1317 (2012).

39 Jadiya, P. et al. Neuronal loss of NCLX-dependent mitochondrial calcium efflux mediates ageassociated cognitive decline. Iscience 26 (2023).

40 Jadiya, P. et al. Impaired mitochondrial calcium efflux contributes to disease progression in models of Alzheimer’s disease. Nature communications 10, 3885 (2019).

41 De Mario, A. et al. Identification and functional validation of FDA-approved positive and negative modulators of the mitochondrial calcium uniporter. Cell Rep 35, 109275 (2021). 10.1016/j.celrep.2021.109275

42 Singh, R. et al. Uncontrolled mitochondrial calcium uptake underlies the pathogenesis of neurodegeneration in MICU1-deficient mice and patients. Science Advances 8, eabj4716 (2022).

43 Logan, C. V. et al. Loss-of-function mutations in MICU1 cause a brain and muscle disorder linked to primary alterations in mitochondrial calcium signaling. Nature genetics 46, 188–193 (2014).

44 Yushkevich, P. A. et al. User-guided 3D active contour segmentation of anatomical structures: significantly improved efficiency and reliability. Neuroimage 31, 1116–1128 (2006). 10.1016/j.neuroimage.2006.01.015

45 Kim, Y., Yang, D. S., Katti, P. & Glancy, B. Protein composition of the muscle mitochondrial reticulum during postnatal development. J Physiol 597, 2707–2727 (2019). 10.1113/JP277579

