## Extended Data Figures for "Loss of TMEM65 causes mitochondrial disease mediated by mitochondrial calcium"

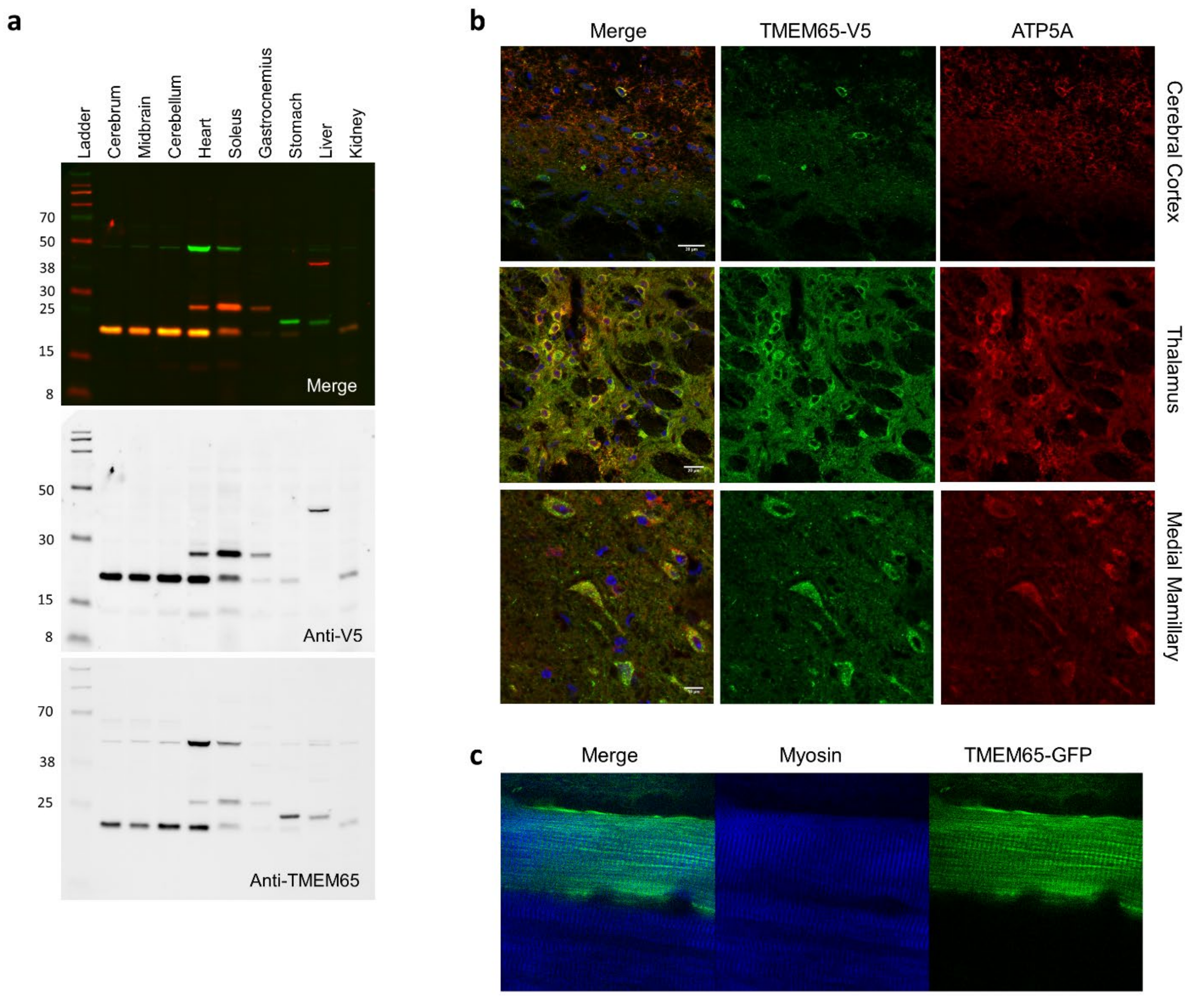


**Extended Data Figure 1. TMEM65 is widely expressed in mouse tissues including brain, heart and skeletal muscles.**

**a**, Representative Western blot analysis of different tissues from an adult *Tmem65*$^{+V5/+V5}$ mouse. Anti-V5 antibody and anti-TMEM65 antibody were used on the same blot to confirm TMEM65 expression in different tissues. **b**, TMEM65 is colocalized with ATP5A, a mitochondrial complex V subunit in mouse brain sections. **c**, mouse TA muscle transfected with TMEM65-GFP plasmid shows grid-like mitochondrial network surrounding contractile machinery of the muscle fiber in vivo.

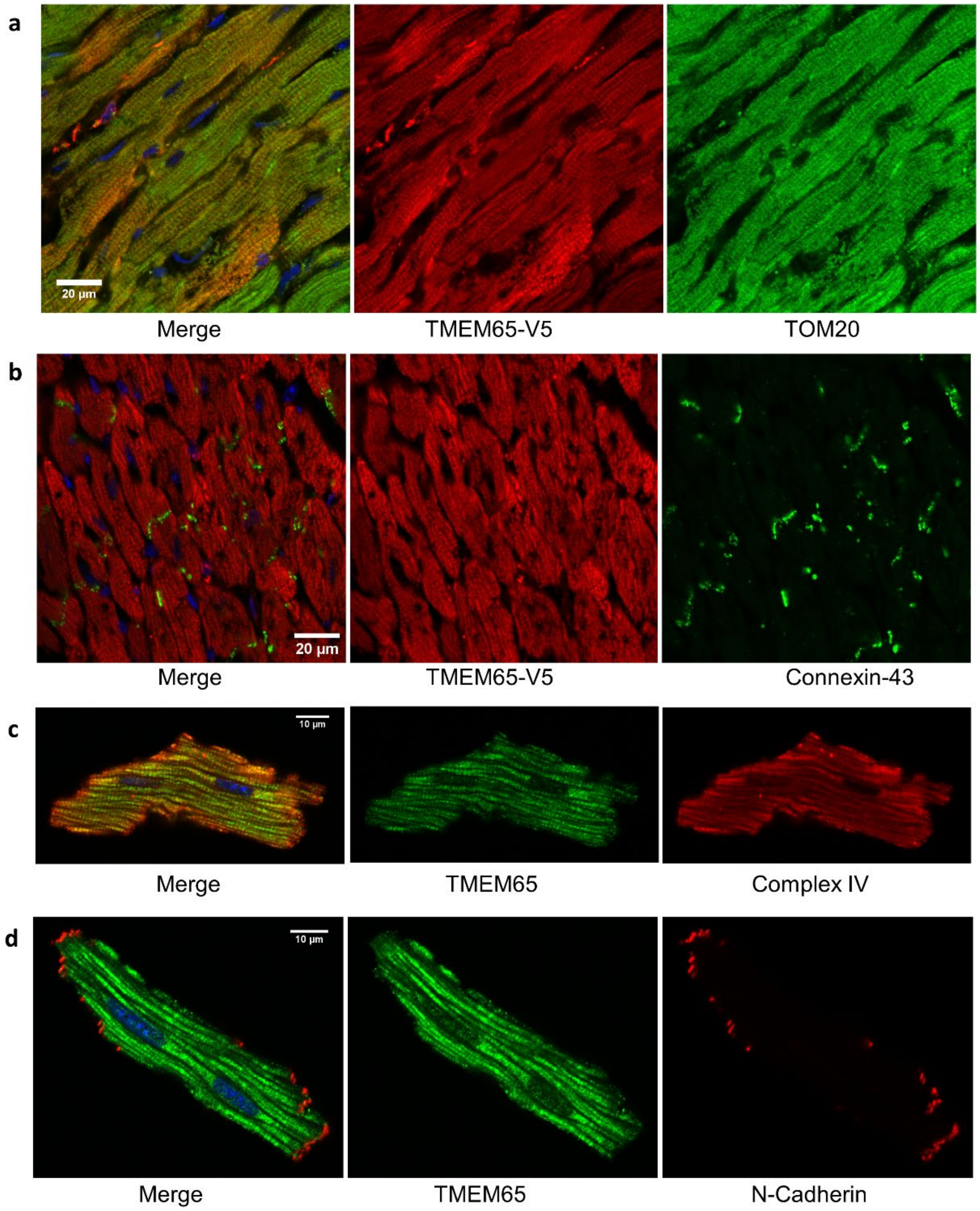


**Extended Data Figure 2. TMEM65 is a mitochondrial protein expressed in mouse heart.**

**a,** Heart section from a *Tmem65*$^{+V5/+V5}$ mouse was stained with anti-V5 and anti-TOM20 antibodies to show TMEM65 is a mitochondrial protein in the heart. **b,** Heart section from a *Tmem65*$^{+V5/+V5}$ mouse was stained with anti-V5 and anti-Connexin-43 antibodies to show TMEM65 is not colocalized with Connexin-43, an intercalated disc protein in mouse heart. **c,** Isolated mouse cardiac myocyte was stained with anti-TMEM65 and anti-Complex IV subunit IV antibodies to show TMEM65 is a mitochondrial protein in cardiac myocyte. **d,** Isolated mouse cardiac myocyte was stained with anti-TMEM65 and anti-N-Cadherin to show that TMEM65 is not colocalized with N-Cadherin, an intercalated disc protein in cardiac myocyte.

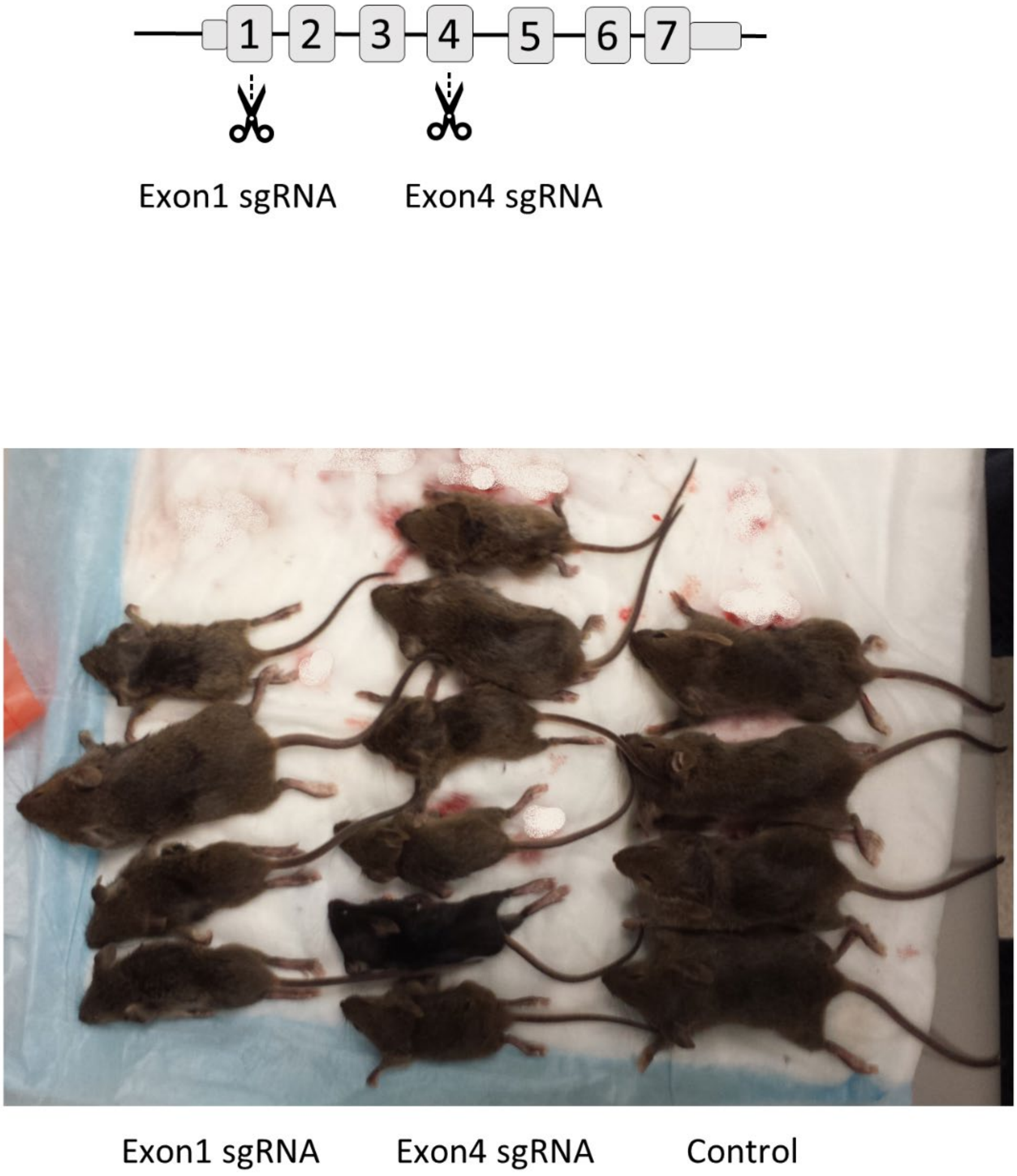


**Extended Data Figure 3. Design of CRISPR global Tmem65 knockout mice.**

Upper: Schematic of global TMEM65 KO mouse with sgRNAs targeting exon1 and/or exon4 of Tmem65 using CRISPR methodology. Lower: Reduced size of global Tmem65 knockout mice with sgRNAs targeting exon1 or exon4.

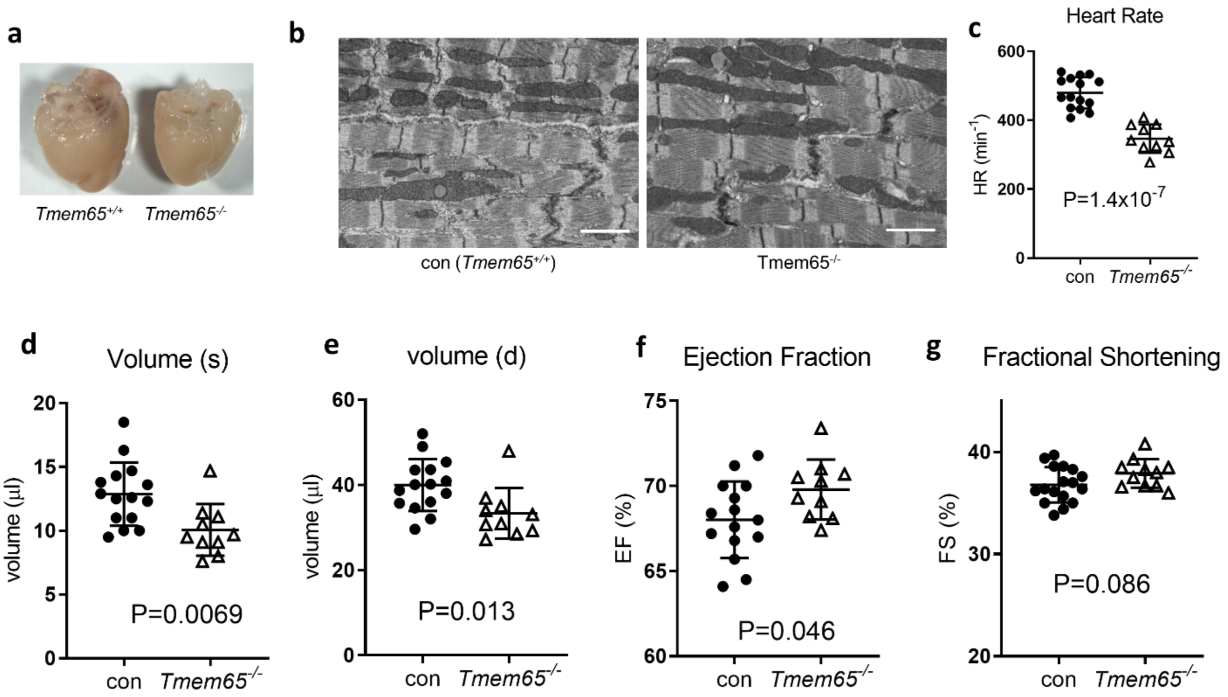


**Extended Data Figure 4. Whole body ablation of TMEM65 does not change heart structure and function.**

**a,** Hearts from P21 littermates of control *Tmem65*$^{+/+}$ and *Tmem65*$^{-/-}$ pups. **b,** EM images of P20 hearts from *Tmem65*$^{+/+}$ control and *Tmem65*$^{-/-}$ littermates. Muscle contractile apparatus, mitochondria and intercalated discs are all normal in *Tmem65*$^{-/-}$ heart. Bar, 1 µm. **c,** *Tmem65*$^{-/-}$ mice have a lower heart rate compared to littermate control of *Tmem65*$^{+/+}$ at P21 **d-g,** Echocardiology results from P21 control *Tmem65*$^{+/+}$ and *Tmem65*$^{-/-}$ littermates. Systolic volumes (**d**) and diastolic volumes (**e**) were significantly lower, while ejection fraction (**f**) was higher in P21 *Tmem65*$^{-/-}$ hearts. **g,** Percentage of fractional shortening was similar between *Tmem65*$^{+/+}$ control and *Tmem65*$^{-/-}$ littermates. *Tmem65*$^{+/+}$ n =15; *Tmem65*$^{-/-}$ n = 10. Individual values as well as mean ± SD are presented.

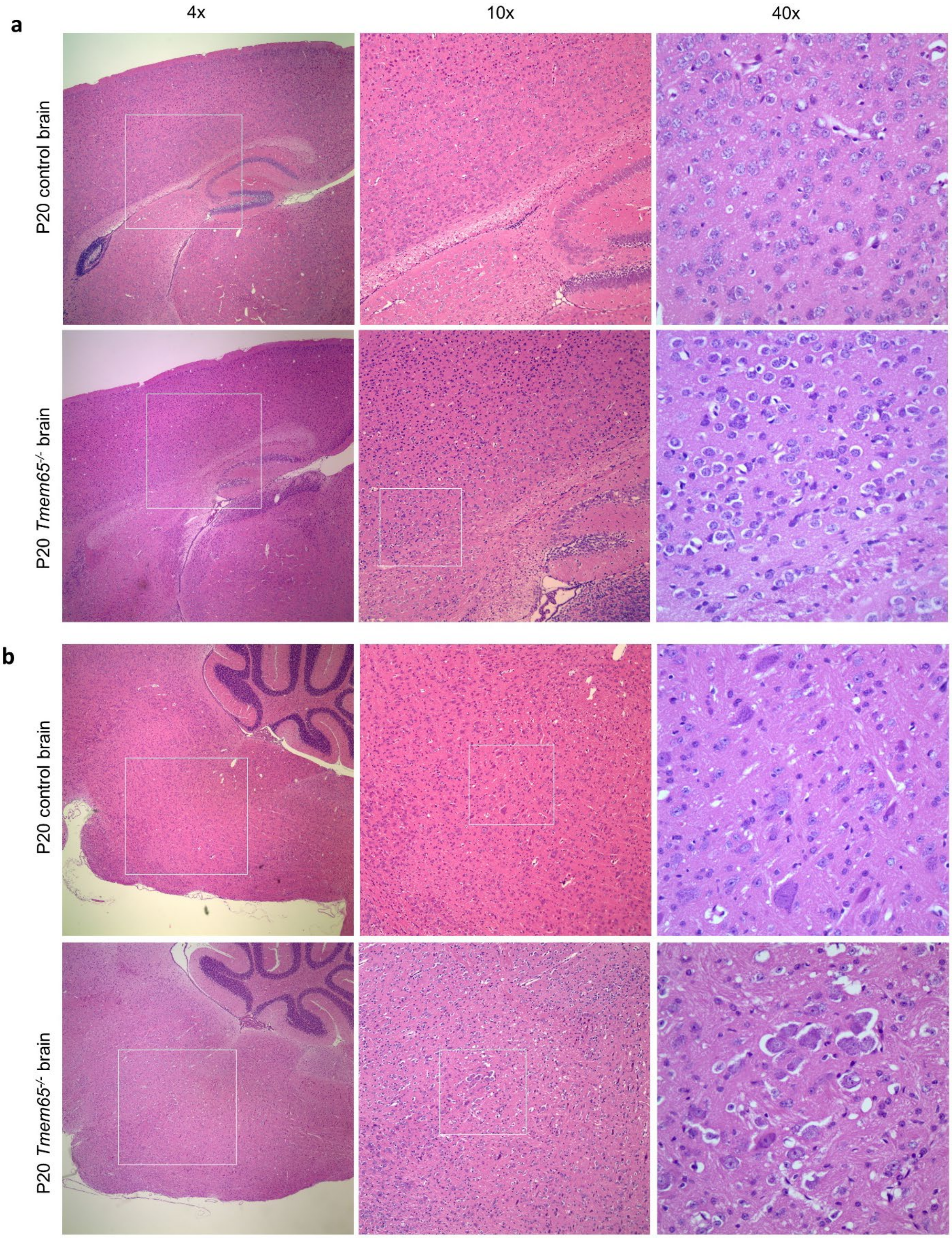


**Extended Data Figure 5. Brain lesions in a P20 *Tmem65*$^{-/-}$ mouse.** H&E staining of brain sagittal sections show diffusive neuronal vacuolar degeneration in the cingulate cortex region (**a**), and brain stem area (**b**). Images were taken at 4x, 10x, and 40x.

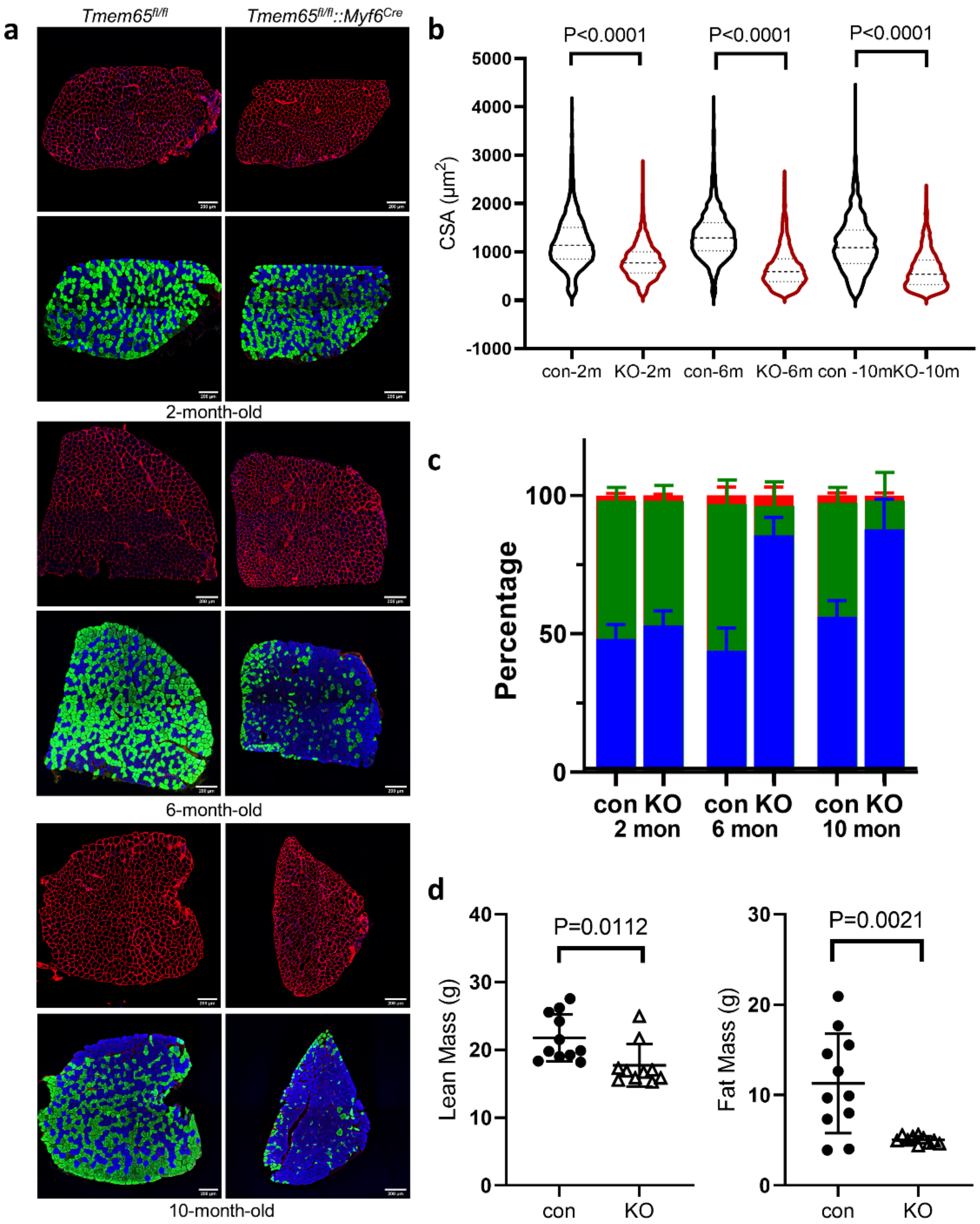


**Extended Data Figure 6. Loss of TMEM65 in skeletal muscles causes muscular atrophy and muscle type switch. a,** Immunofluorescence of soleus muscles from control *Tmem65*$^{fl/fl}$ and *Tmem65*$^{fl/fl}$ *::Myf6*$^{Cre}$ littermates at 2-month-old, 6-month-old, and 10-month-old. Anti-Laminin antibody staining (red, upper rows) delineated individual muscle fibers for cross-section area quantification and evaluation. Myosin isoforms were identified with specific antibodies to muscle type I (blue), IIa (green) and IIb (red), in lower rows. **b,** Quantification of soleus muscle fiber cross section area from control *Tmem65*$^{fl/fl}$ and *Tmem65*$^{fl/fl}$ *::Myf6*$^{Cre}$ littermates of different ages. **c,** Quantification of myosin isoform composition in control *Tmem65*$^{fl/fl}$ and *Tmem65*$^{fl/fl}$ *::Myf6*$^{Cre}$ soleus muscle cross sections at different age.(n = 3 replicates per group). **d,** EchoMRI body composition analysis of *Tmem65*$^{fl/fl}$*::Myf6*$^{Cre}$ mouse at 6-month-old. *Tmem65*$^{fl/fl}$ n = 11; *Tmem65*$^{fl/fl}$*::Myf6*$^{Cre}$ n = 10. Individual values as well as mean ± SD are presented.

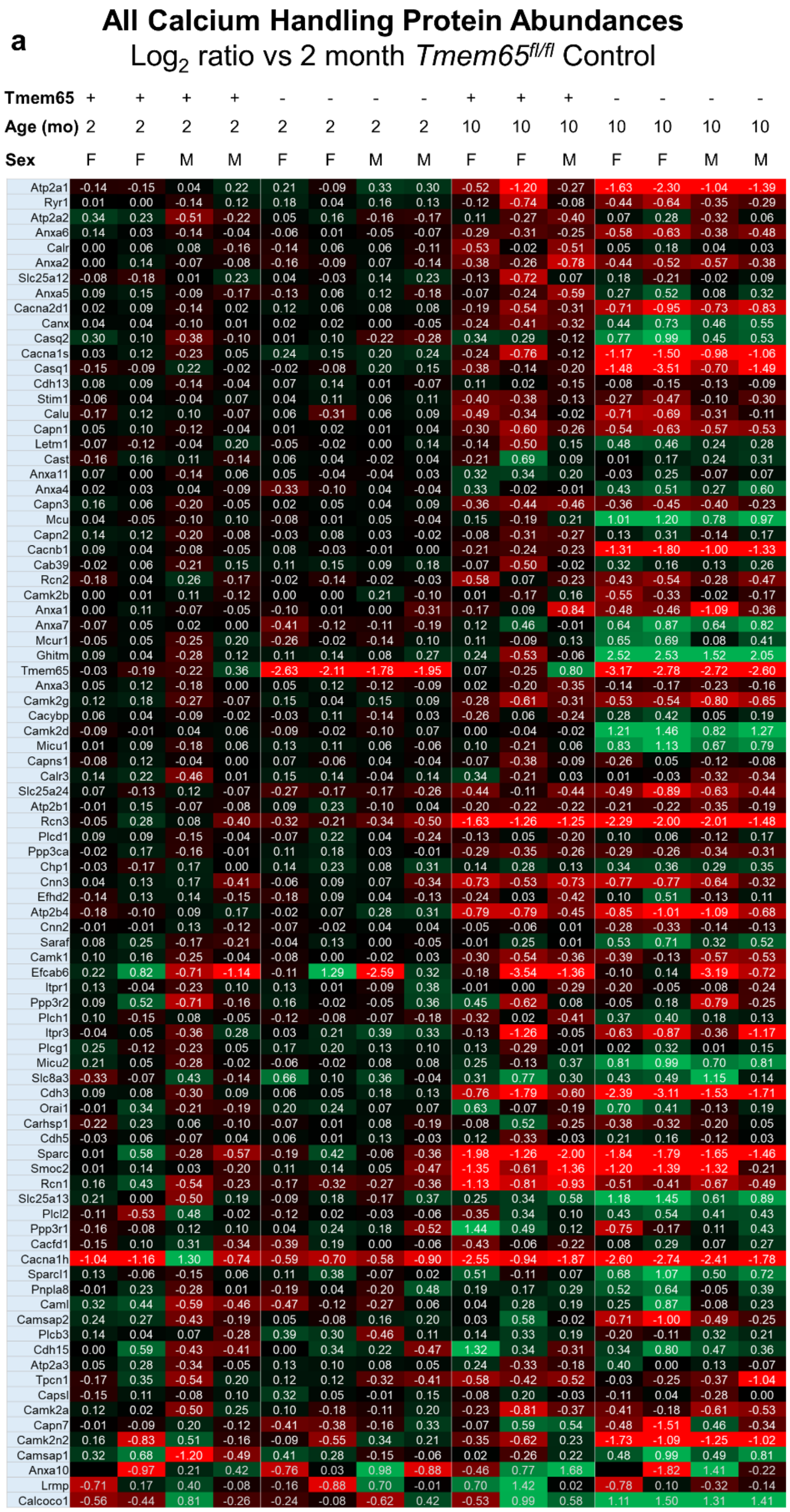


| Tmem65 | + | + | + | + | - | - | - | - | + | + | + | - | - | - | - |
|---|---|---|---|---|---|---|---|---|---|---|---|---|---|---|---|
| Age (mo) | 2 | 2 | 2 | 2 | 2 | 2 | 2 | 2 | 10 | 10 | 10 | 10 | 10 | 10 | 10 |
| Sex | F | F | M | M | F | F | M | M | F | F | M | F | F | M | M |
| Atp2a1 | -0.14 | -0.15 | 0.04 | 0.22 | 0.21 | -0.09 | 0.33 | 0.30 | -0.52 | -1.20 | -0.27 | -1.63 | -2.30 | -1.04 | -1.39 |
| Ryr1 | 0.01 | 0.00 | -0.14 | 0.12 | 0.18 | 0.04 | 0.16 | 0.13 | -0.12 | -0.74 | -0.08 | -0.44 | -0.64 | -0.35 | -0.29 |
| Atp2a2 | 0.34 | 0.23 | -0.51 | -0.22 | 0.05 | 0.16 | -0.16 | -0.17 | 0.11 | -0.27 | -0.40 | 0.07 | 0.28 | -0.32 | 0.06 |
| Anxa6 | 0.14 | 0.03 | -0.14 | -0.04 | -0.06 | 0.01 | -0.05 | -0.07 | -0.29 | -0.31 | -0.25 | -0.58 | -0.63 | -0.38 | -0.48 |
| Calr | 0.00 | 0.06 | 0.08 | -0.16 | -0.14 | 0.06 | 0.06 | -0.11 | -0.53 | -0.02 | -0.51 | 0.05 | 0.18 | 0.04 | 0.03 |
| Anxa2 | 0.00 | 0.14 | -0.07 | -0.08 | -0.16 | -0.09 | 0.07 | -0.14 | -0.38 | -0.26 | -0.78 | -0.44 | -0.52 | -0.57 | -0.38 |
| Slc25a12 | -0.08 | -0.18 | 0.01 | 0.23 | 0.04 | -0.03 | 0.14 | 0.23 | -0.13 | -0.72 | 0.07 | 0.18 | -0.21 | -0.02 | 0.09 |
| Anxa5 | 0.09 | 0.15 | -0.09 | -0.17 | -0.13 | 0.06 | 0.12 | -0.18 | -0.07 | -0.24 | -0.59 | 0.27 | 0.52 | 0.08 | 0.32 |
| Cacna2d1 | 0.02 | 0.09 | -0.14 | 0.02 | 0.12 | 0.06 | 0.08 | 0.08 | -0.19 | -0.54 | -0.31 | -0.71 | -0.95 | -0.73 | -0.83 |
| Canx | 0.04 | 0.04 | -0.10 | 0.01 | 0.02 | 0.02 | 0.00 | -0.05 | -0.24 | -0.41 | -0.32 | 0.44 | 0.73 | 0.46 | 0.55 |
| Casq2 | 0.30 | 0.10 | -0.38 | -0.10 | 0.01 | 0.10 | -0.22 | -0.28 | 0.34 | 0.29 | -0.12 | 0.77 | 0.99 | 0.45 | 0.53 |
| Cacna1s | 0.03 | 0.12 | -0.23 | 0.05 | 0.24 | 0.15 | 0.20 | 0.24 | -0.24 | -0.76 | -0.12 | -1.17 | -1.50 | -0.98 | -1.06 |
| Casq1 | -0.15 | -0.09 | 0.22 | -0.02 | -0.02 | -0.08 | 0.20 | 0.15 | -0.38 | -0.14 | -0.20 | -1.48 | -3.51 | -0.70 | -1.49 |
| Cdh13 | 0.08 | 0.09 | -0.14 | -0.04 | 0.07 | 0.14 | 0.01 | -0.07 | 0.11 | 0.02 | -0.15 | -0.08 | -0.15 | -0.13 | -0.09 |
| Stim1 | -0.06 | 0.04 | -0.04 | 0.07 | 0.04 | 0.11 | 0.06 | 0.11 | -0.40 | -0.38 | -0.13 | -0.27 | -0.47 | -0.10 | -0.30 |
| Calu | -0.17 | 0.12 | 0.10 | -0.07 | 0.06 | -0.31 | 0.06 | 0.09 | -0.49 | -0.34 | -0.02 | -0.71 | -0.69 | -0.31 | -0.11 |
| Capn1 | 0.05 | 0.10 | -0.12 | -0.04 | 0.01 | 0.02 | 0.01 | 0.04 | -0.30 | -0.60 | -0.26 | -0.54 | -0.63 | -0.57 | -0.53 |
| Letm1 | -0.07 | -0.12 | -0.04 | 0.20 | -0.05 | -0.02 | 0.00 | 0.14 | -0.14 | -0.50 | 0.15 | 0.48 | 0.46 | 0.24 | 0.28 |
| Cast | -0.16 | 0.16 | 0.11 | -0.14 | 0.06 | 0.04 | -0.02 | 0.04 | -0.21 | 0.69 | 0.09 | 0.01 | 0.17 | 0.24 | 0.31 |
| Anxa11 | 0.07 | 0.00 | -0.14 | 0.06 | 0.05 | -0.04 | -0.04 | 0.03 | 0.32 | 0.34 | 0.20 | -0.03 | 0.25 | -0.07 | 0.07 |
| Anxa4 | 0.02 | 0.03 | 0.04 | -0.09 | -0.33 | -0.10 | 0.04 | -0.04 | 0.33 | -0.02 | -0.01 | 0.43 | 0.51 | 0.27 | 0.60 |
| Capn3 | 0.16 | 0.06 | -0.20 | -0.05 | 0.02 | 0.05 | 0.04 | 0.09 | -0.36 | -0.44 | -0.46 | -0.36 | -0.45 | -0.40 | -0.23 |
| Mcu | 0.04 | -0.05 | -0.10 | 0.10 | -0.08 | 0.01 | 0.05 | -0.04 | 0.15 | -0.19 | 0.21 | 1.01 | 1.20 | 0.78 | 0.97 |
| Capn2 | 0.14 | 0.12 | -0.20 | -0.08 | -0.03 | 0.08 | 0.03 | -0.02 | -0.08 | -0.31 | -0.27 | 0.13 | 0.31 | -0.14 | 0.17 |
| Cacnb1 | 0.09 | 0.04 | -0.08 | -0.05 | 0.08 | -0.03 | -0.01 | 0.00 | -0.21 | -0.24 | -0.23 | -1.31 | -1.80 | -1.00 | -1.33 |
| Cab39 | -0.02 | 0.06 | -0.21 | 0.15 | 0.11 | 0.15 | 0.09 | 0.18 | -0.07 | -0.50 | -0.02 | 0.32 | 0.16 | 0.13 | 0.26 |
| Rcn2 | -0.18 | 0.04 | 0.26 | -0.17 | -0.02 | -0.14 | -0.02 | -0.03 | -0.58 | 0.07 | -0.23 | -0.43 | -0.54 | -0.28 | -0.47 |
| Camk2b | 0.00 | 0.01 | 0.11 | -0.12 | 0.00 | 0.00 | 0.21 | -0.10 | 0.01 | -0.17 | 0.16 | -0.55 | -0.33 | -0.02 | -0.17 |
| Anxa1 | 0.00 | 0.11 | -0.07 | -0.05 | -0.10 | 0.01 | 0.00 | -0.31 | -0.17 | 0.09 | -0.84 | -0.48 | -0.46 | -1.09 | -0.36 |
| Anxa7 | -0.07 | 0.05 | 0.02 | 0.00 | -0.41 | -0.12 | -0.11 | -0.19 | 0.12 | 0.46 | -0.01 | 0.64 | 0.87 | 0.64 | 0.82 |
| Mcur1 | -0.05 | 0.05 | -0.25 | 0.20 | -0.26 | -0.02 | -0.14 | 0.10 | 0.11 | -0.09 | 0.13 | 0.65 | 0.69 | 0.08 | 0.41 |
| Ghitm | 0.09 | 0.04 | -0.28 | 0.12 | 0.11 | 0.14 | 0.08 | 0.27 | 0.24 | -0.53 | -0.06 | 2.52 | 2.53 | 1.52 | 2.05 |
| Tmem65 | -0.03 | -0.19 | -0.22 | 0.36 | -2.63 | -2.11 | -1.78 | -1.95 | 0.07 | -0.25 | 0.80 | -3.17 | -2.78 | -2.72 | -2.60 |
| Anxa3 | 0.05 | 0.12 | -0.18 | 0.00 | 0.05 | 0.12 | -0.12 | -0.09 | 0.02 | -0.20 | -0.35 | -0.14 | -0.17 | -0.23 | -0.16 |
| Camk2g | 0.12 | 0.18 | -0.27 | -0.07 | 0.15 | 0.04 | 0.15 | 0.09 | -0.28 | -0.61 | -0.31 | -0.53 | -0.54 | -0.80 | -0.65 |
| Cacybp | 0.06 | 0.04 | -0.09 | -0.02 | -0.03 | 0.11 | -0.14 | 0.03 | -0.26 | 0.06 | -0.24 | 0.28 | 0.42 | 0.05 | 0.19 |
| Camk2d | -0.09 | -0.01 | 0.04 | 0.06 | -0.09 | -0.02 | -0.10 | 0.07 | 0.00 | -0.04 | -0.02 | 1.21 | 1.46 | 0.82 | 1.27 |
| Micu1 | 0.01 | 0.09 | -0.18 | 0.06 | 0.13 | 0.11 | 0.06 | -0.06 | 0.10 | -0.21 | 0.06 | 0.83 | 1.13 | 0.67 | 0.79 |
| Capns1 | -0.08 | 0.12 | -0.04 | 0.00 | 0.07 | -0.06 | 0.04 | -0.04 | -0.07 | -0.38 | -0.09 | -0.26 | 0.05 | -0.12 | -0.08 |
| Calr3 | 0.14 | 0.22 | -0.46 | 0.01 | 0.15 | 0.14 | -0.04 | 0.14 | 0.34 | -0.21 | 0.03 | 0.01 | -0.03 | -0.32 | -0.34 |
| Slc25a24 | 0.07 | -0.13 | 0.12 | -0.07 | -0.27 | -0.17 | -0.17 | -0.26 | -0.44 | -0.11 | -0.44 | -0.49 | -0.89 | -0.63 | -0.44 |
| Atp2b1 | -0.01 | 0.15 | -0.07 | -0.08 | 0.09 | 0.23 | -0.10 | 0.04 | -0.20 | -0.22 | -0.22 | -0.21 | -0.22 | -0.35 | -0.19 |
| Rcn3 | -0.05 | 0.28 | 0.08 | -0.40 | -0.32 | -0.21 | -0.34 | -0.50 | -1.63 | -1.26 | -1.25 | -2.29 | -2.00 | -2.01 | -1.48 |
| Plcd1 | 0.09 | 0.09 | -0.15 | -0.04 | -0.07 | 0.22 | 0.04 | -0.24 | -0.13 | 0.05 | -0.20 | 0.10 | 0.06 | -0.12 | 0.17 |
| Ppp3ca | -0.02 | 0.17 | -0.16 | -0.01 | 0.11 | 0.18 | 0.03 | -0.01 | -0.29 | -0.35 | -0.26 | -0.29 | -0.26 | -0.34 | -0.31 |
| Chp1 | -0.03 | -0.17 | 0.17 | 0.00 | 0.14 | 0.23 | 0.08 | 0.31 | 0.14 | 0.28 | 0.13 | 0.34 | 0.36 | 0.29 | 0.35 |
| Cnn3 | 0.04 | 0.13 | 0.17 | -0.41 | -0.06 | 0.09 | 0.07 | -0.34 | -0.73 | -0.53 | -0.73 | -0.77 | -0.77 | -0.64 | -0.32 |
| Efhd2 | -0.14 | 0.13 | 0.14 | -0.15 | -0.18 | 0.09 | 0.04 | -0.13 | -0.24 | 0.03 | -0.42 | 0.10 | 0.51 | -0.13 | 0.11 |
| Atp2b4 | -0.18 | -0.10 | 0.09 | 0.17 | -0.02 | 0.07 | 0.28 | 0.31 | -0.79 | -0.79 | -0.45 | -0.85 | -1.01 | -1.09 | -0.68 |
| Cnn2 | -0.01 | -0.01 | 0.13 | -0.12 | -0.07 | -0.02 | 0.04 | 0.04 | -0.05 | -0.06 | 0.01 | -0.28 | -0.33 | -0.14 | -0.13 |
| Saraf | 0.08 | 0.25 | -0.17 | -0.21 | -0.04 | 0.13 | 0.00 | -0.05 | -0.01 | 0.25 | 0.01 | 0.53 | 0.71 | 0.32 | 0.52 |
| Camk1 | 0.10 | 0.16 | -0.25 | -0.04 | -0.08 | 0.00 | -0.02 | 0.03 | -0.30 | -0.54 | -0.36 | -0.39 | -0.13 | -0.57 | -0.53 |
| Efcab6 | 0.22 | 0.82 | -0.71 | -1.14 | -0.11 | 1.29 | -2.59 | 0.32 | -0.18 | -3.54 | -1.36 | -0.10 | 0.14 | -3.19 | -0.72 |
| Itpr1 | 0.13 | -0.04 | -0.23 | 0.10 | 0.13 | 0.01 | -0.09 | 0.38 | -0.01 | 0.00 | -0.29 | -0.20 | -0.05 | -0.08 | -0.24 |
| Ppp3r2 | 0.09 | 0.52 | -0.71 | -0.16 | 0.16 | -0.02 | -0.05 | 0.36 | 0.45 | -0.62 | 0.08 | -0.05 | 0.18 | -0.79 | -0.25 |
| Plch1 | 0.10 | -0.15 | 0.08 | -0.05 | -0.12 | -0.08 | -0.07 | -0.18 | -0.32 | 0.02 | -0.41 | 0.37 | 0.40 | 0.18 | 0.13 |
| Itpr3 | -0.04 | 0.05 | -0.36 | 0.28 | 0.03 | 0.21 | 0.39 | 0.33 | -0.13 | -1.26 | -0.05 | -0.63 | -0.87 | -0.36 | -1.17 |
| Plcg1 | 0.25 | -0.12 | -0.23 | 0.05 | 0.17 | 0.20 | 0.13 | 0.10 | 0.13 | -0.29 | -0.01 | 0.02 | 0.32 | 0.01 | 0.15 |
| Micu2 | 0.21 | 0.05 | -0.28 | -0.02 | -0.06 | -0.02 | 0.08 | 0.08 | 0.25 | -0.13 | 0.37 | 0.81 | 0.99 | 0.70 | 0.81 |
| Slc8a3 | -0.33 | -0.07 | 0.43 | -0.14 | 0.66 | 0.10 | 0.36 | -0.04 | 0.31 | 0.77 | 0.30 | 0.43 | 0.49 | 1.15 | 0.14 |
| Cdh3 | 0.09 | 0.08 | -0.30 | 0.09 | 0.06 | 0.05 | 0.18 | 0.13 | -0.76 | -1.79 | -0.60 | -2.39 | -3.11 | -1.53 | -1.71 |
| Orai1 | -0.01 | 0.34 | -0.21 | -0.19 | 0.20 | 0.24 | 0.07 | 0.07 | 0.63 | -0.07 | -0.19 | 0.70 | 0.41 | -0.13 | 0.19 |
| Carhsp1 | -0.22 | 0.23 | 0.06 | -0.10 | -0.07 | 0.01 | 0.08 | -0.19 | -0.08 | 0.52 | -0.25 | -0.38 | -0.32 | -0.20 | 0.05 |
| Cdh5 | -0.03 | 0.06 | -0.07 | 0.04 | 0.06 | 0.01 | 0.13 | -0.03 | 0.12 | -0.33 | -0.03 | 0.21 | 0.16 | -0.12 | 0.03 |
| Sparc | 0.01 | 0.58 | -0.28 | -0.57 | -0.19 | 0.42 | -0.06 | -0.36 | -1.98 | -1.26 | -2.00 | -1.84 | -1.79 | -1.65 | -1.46 |
| Smoc2 | 0.01 | 0.14 | 0.03 | -0.20 | 0.11 | 0.14 | 0.05 | -0.47 | -1.35 | -0.61 | -1.36 | -1.20 | -1.39 | -1.32 | -0.21 |
| Rcn1 | 0.16 | 0.43 | -0.54 | -0.23 | -0.17 | -0.32 | -0.27 | -0.36 | -1.13 | -0.81 | -0.93 | -0.51 | -0.41 | -0.67 | -0.49 |
| Slc25a13 | 0.21 | 0.00 | -0.50 | 0.19 | -0.09 | 0.18 | -0.17 | 0.37 | 0.25 | 0.34 | 0.58 | 1.18 | 1.45 | 0.61 | 0.89 |
| Plcl2 | -0.11 | -0.53 | 0.48 | -0.02 | -0.12 | 0.02 | -0.03 | -0.06 | -0.35 | 0.34 | 0.10 | 0.43 | 0.54 | 0.41 | 0.43 |
| Ppp3r1 | -0.16 | -0.08 | 0.12 | 0.10 | 0.04 | 0.24 | 0.18 | -0.52 | 1.44 | 0.49 | 0.12 | -0.75 | -0.17 | 0.11 | 0.43 |
| Cacfd1 | -0.15 | 0.10 | 0.31 | -0.34 | -0.39 | 0.19 | 0.00 | -0.06 | -0.43 | -0.06 | -0.22 | 0.08 | 0.29 | 0.07 | 0.27 |
| Cacna1h | -1.04 | -1.16 | 1.30 | -0.74 | -0.59 | -0.70 | -0.58 | -0.90 | -2.55 | -0.94 | -1.87 | -2.60 | -2.74 | -2.41 | -1.78 |
| Sparcl1 | 0.13 | -0.06 | -0.15 | 0.06 | 0.11 | 0.38 | -0.07 | 0.02 | 1.32 | -0.11 | 0.07 | 0.68 | 1.07 | 0.50 | 0.72 |
| Pnpla8 | -0.01 | 0.23 | -0.28 | 0.01 | -0.19 | 0.04 | -0.20 | 0.48 | 0.19 | 0.17 | 0.29 | 0.52 | 0.64 | -0.05 | 0.39 |
| Caml | 0.32 | 0.44 | -0.59 | -0.46 | -0.47 | -0.12 | -0.27 | 0.06 | 0.04 | 0.28 | 0.19 | 0.25 | 0.87 | -0.08 | 0.23 |
| Camsap2 | 0.24 | 0.27 | -0.43 | -0.19 | 0.05 | -0.08 | 0.16 | 0.20 | 0.03 | 0.58 | -0.02 | -0.71 | -1.00 | -0.49 | -0.25 |
| Plcb3 | 0.14 | 0.04 | 0.07 | -0.28 | 0.39 | 0.30 | -0.46 | 0.11 | 0.14 | 0.33 | 0.19 | -0.20 | -0.11 | 0.32 | 0.21 |
| Cdh15 | 0.00 | 0.59 | -0.43 | -0.41 | 0.00 | 0.34 | 0.22 | -0.47 | 0.51 | 0.34 | -0.31 | 0.34 | 0.80 | 0.47 | 0.36 |
| Atp2a3 | 0.05 | 0.28 | -0.34 | -0.05 | 0.13 | 0.10 | 0.08 | 0.05 | 0.24 | -0.33 | -0.18 | 0.40 | 0.00 | 0.13 | -0.07 |
| Tpcn1 | -0.17 | 0.35 | -0.54 | 0.20 | 0.12 | 0.12 | -0.32 | -0.41 | -0.58 | -0.42 | -0.52 | -0.03 | -0.25 | -0.37 | -1.04 |
| Capsl | -0.15 | 0.11 | -0.08 | 0.10 | 0.32 | 0.05 | -0.01 | 0.15 | -0.08 | 0.20 | -0.03 | -0.11 | 0.04 | -0.28 | 0.00 |
| Camk2a | 0.12 | 0.02 | -0.50 | 0.25 | 0.10 | -0.18 | -0.11 | 0.20 | -0.23 | -0.81 | -0.37 | -0.41 | -0.18 | -0.61 | -0.53 |
| Capn7 | -0.01 | -0.09 | 0.20 | -0.12 | -0.41 | -0.38 | -0.16 | 0.33 | -0.07 | 0.59 | 0.54 | -0.48 | -1.51 | 0.46 | -0.34 |
| Camk2n2 | 0.16 | -0.83 | 0.51 | -0.16 | -0.09 | -0.55 | 0.34 | 0.21 | -0.35 | -0.62 | 0.23 | -1.73 | -1.09 | -1.25 | -1.02 |
| Camsap1 | 0.32 | 0.68 | -1.20 | -0.49 | 0.41 | 0.28 | -0.15 | -0.06 | 0.00 | -0.26 | 0.22 | 0.48 | 0.99 | 0.49 | 0.81 |
| Anxa10 | | -0.97 | 0.21 | 0.42 | -0.76 | 0.03 | 0.98 | -0.88 | -0.46 | 0.77 | 1.68 | | -1.82 | 1.41 | -0.22 |
| Lrmp | -0.71 | 0.17 | 0.40 | -0.08 | -0.16 | -0.88 | 0.70 | -0.01 | 0.70 | 1.42 | 0.02 | -0.78 | 0.10 | -0.32 | -0.14 |
| Calcoco1 | -0.56 | -0.44 | 0.81 | -0.26 | -0.24 | -0.08 | -0.62 | 0.42 | -0.53 | 0.99 | 0.58 | 1.11 | 1.50 | 1.31 | 1.41 |

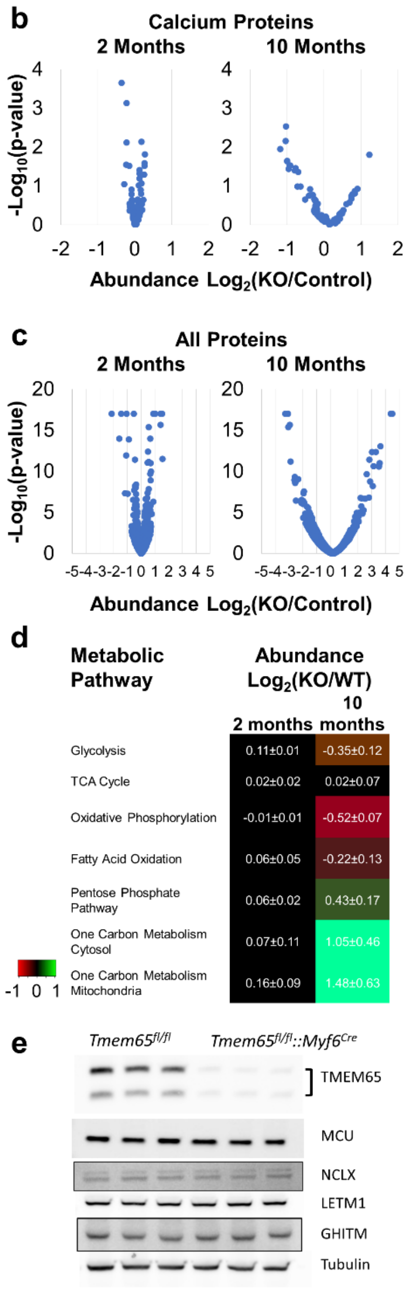


**Extended Data Figure 7. Proteomic analysis of muscle-specific TMEM65 KO and control muscles.** **a,** Abundances of all calcium handling proteins at 2 months and 10 months of age in TMEM65 KO and control muscles relative to mean 2-month control values. Proteins sorted top to bottom by abundance. **b,** Volcano plot of 2 month calcium handling protein abundances from **a**. **c,** Volcano plot of 10 month protein abundances from **a**. **d,** Abundances of proteins within major metabolic pathways at 2 months and 10 months. **e,** Western blot analysis of 2-month-old soleus from *Tmem65$^{fl/fl}$* or *Tmem65$^{fl/fl}$::Myf6$^{Cre}$* mice. n = 3 replicates per group.

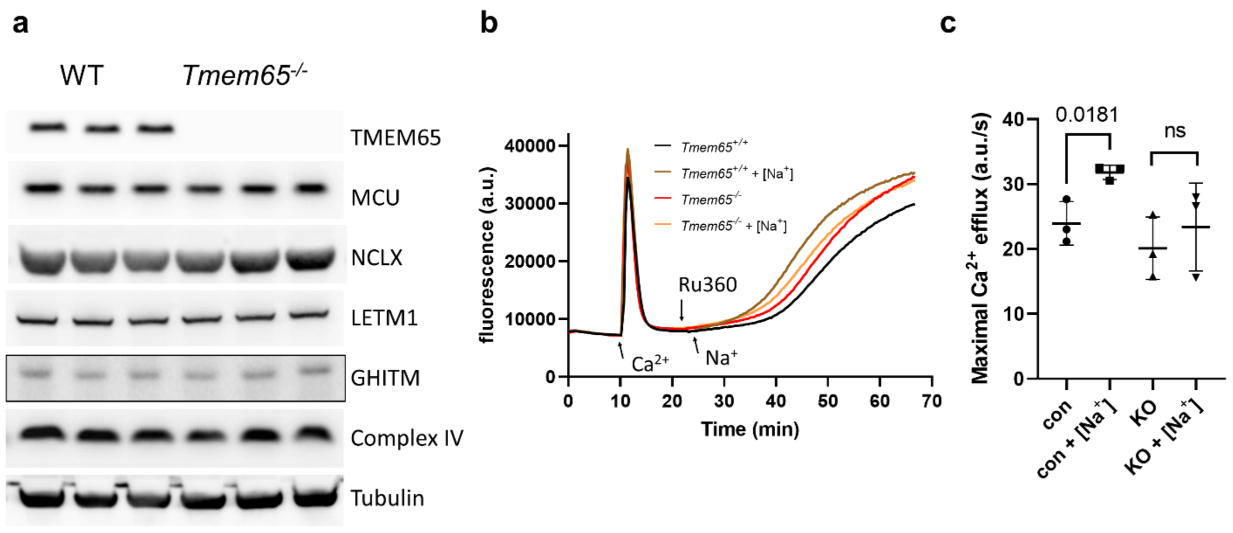


**Extended Data Figure 8. TMEM65 regulates brain mitochondrial calcium release in $Na^+$ dependent manner**

**a,** Western blot analysis of TMEM65, MCU, NCLX, LETM1, and GHITM protein expression in brain tissues (n = 3 replicates per group). **b,** Representative traces of extramitochondrial $Ca^{2+}$ ($[Ca^{2+}]_{ext}$) in a suspension of isolated brain mitochondria (100 µg/well). First, a bolus of $Ca^{2+}$ (50 µM) was added, followed by addition of MCU inhibitor, Ru360 (3 µM) to block mitochondrial $Ca^{2+}$ uptake; then 20 mM NaCl was added to induce $Ca^{2+}$ from mitochondria. **c,** Summary of the maximal rates of mitochondrial $Ca^{2+}$ efflux induced by Ru360 alone, or Ru360 and $Na^+$ in *Tmem65*$^{fl/fl}$ control or *Tmem65*$^{fl/fl}$*::CMV*$^{Cre}$ mitochondria isolated from brain. n = 3 replicates per group.
