## Supplemental Figures for "Loss of TMEM65 causes mitochondrial disease mediated by mitochondrial calcium"

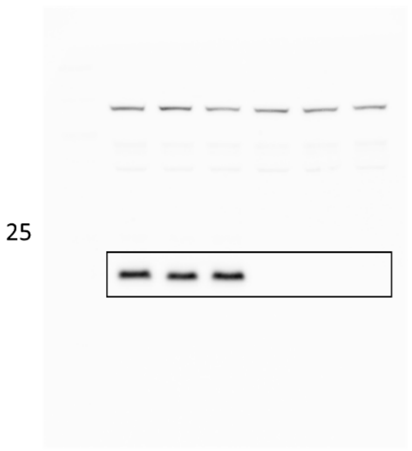


Brain

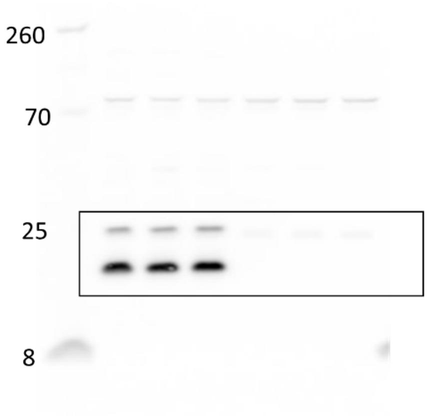


Heart

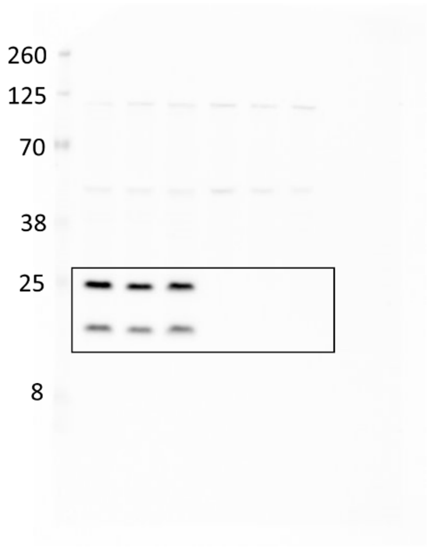


Skeletal Muscle

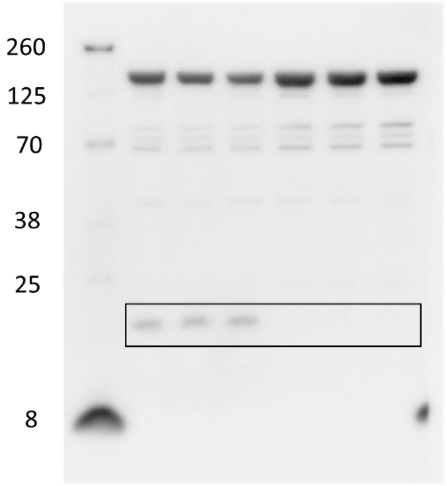


Liver

Western blots of Figure 1d with anti-TMEM65 antibody.

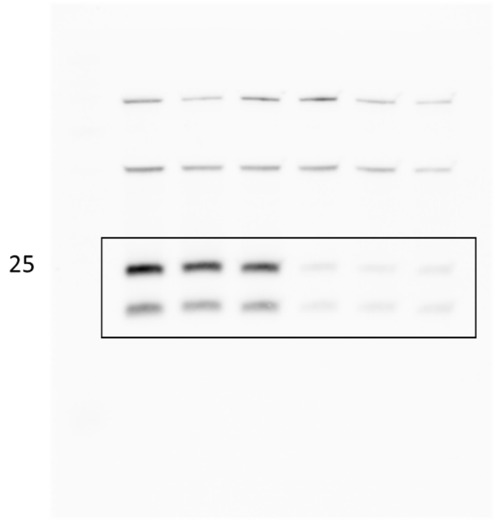


TMEM65

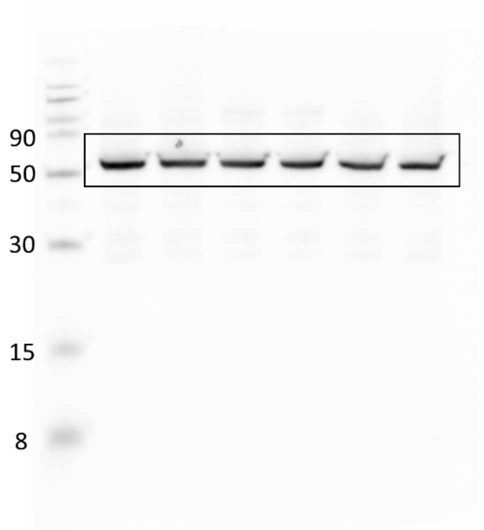


Tubulin

Western blots of Figure 3a

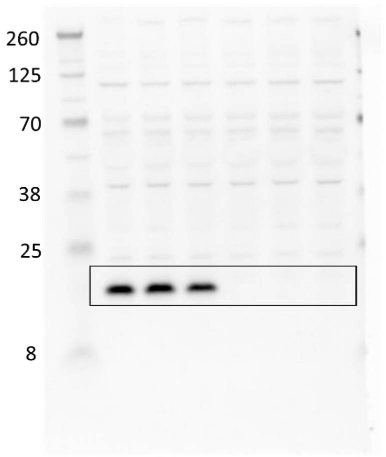


TMEM65

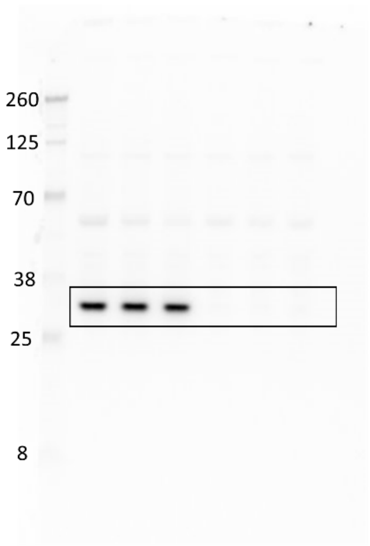


MCU

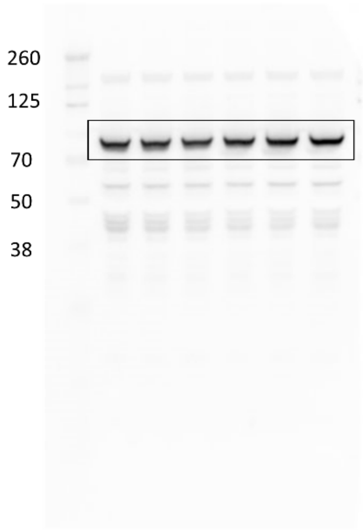


LETM1

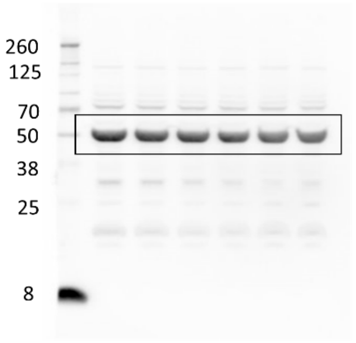


NCLX

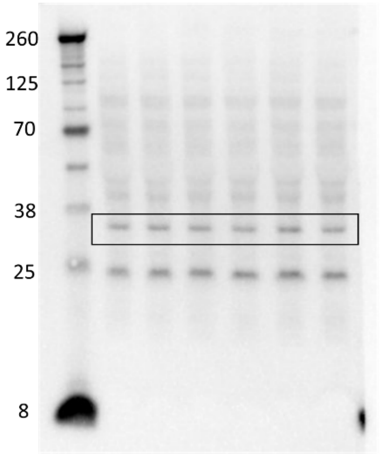


GHITM

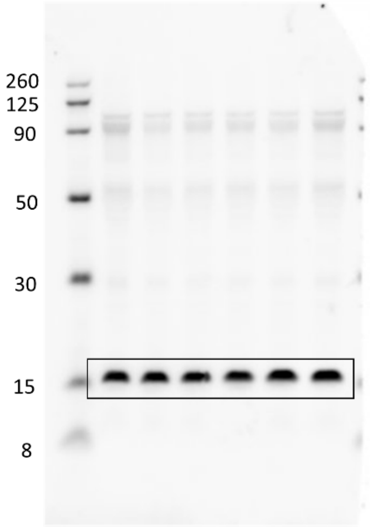


Complex IV subunit IV

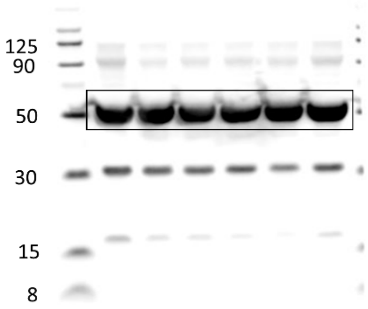


Tubulin

Western blots of Figure 4g.

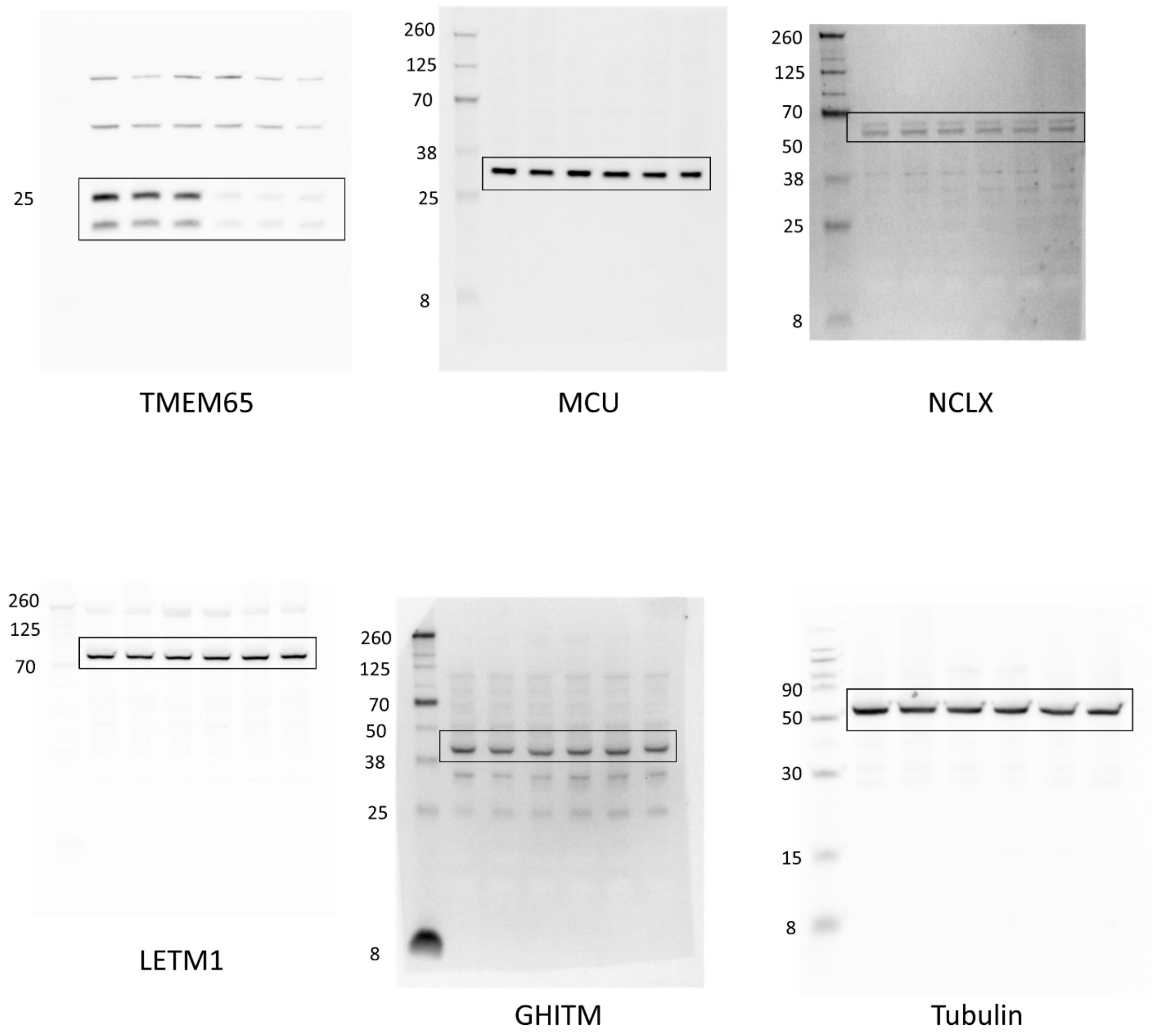


Western blots of Extended Data Figure 7e

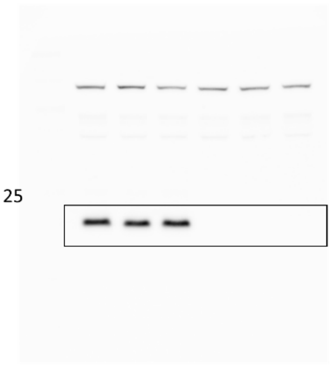


TMEM65

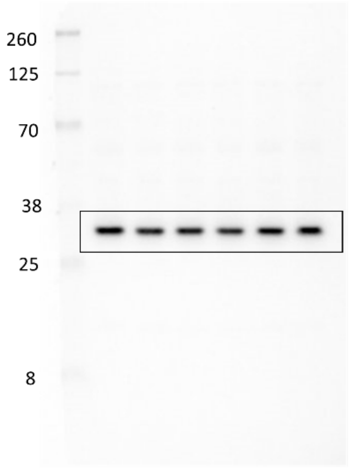


MCU

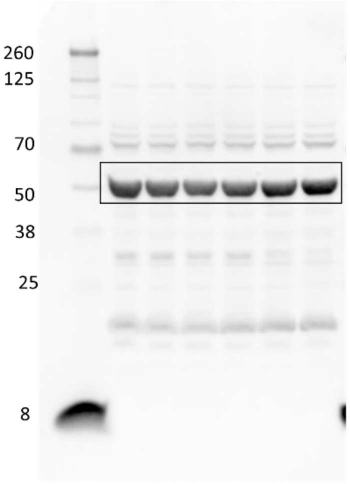


NCLX

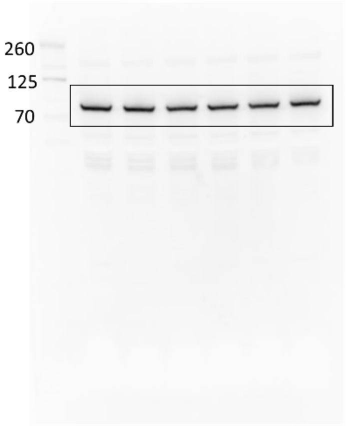


LETM1

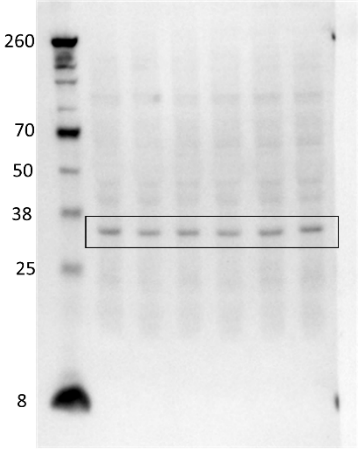


GHITM

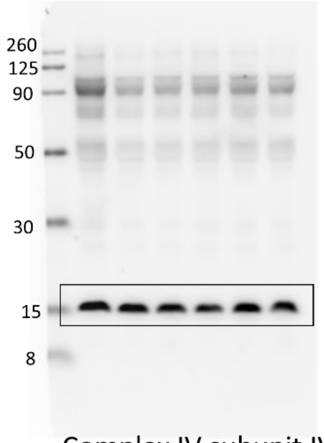


Complex IV subunit IV

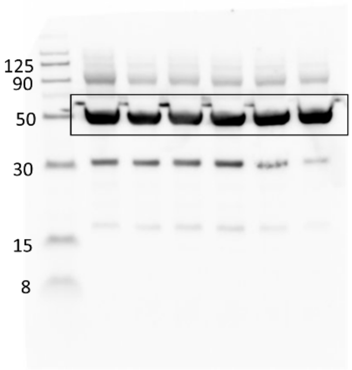


Tubulin

Western blots of Extended Data Figure 8a
